# Integrated inference of cancer gene expression from cell-free plasma chromatin

**DOI:** 10.64898/2026.02.18.706026

**Authors:** Gunsagar S. Gulati, Damien Vasseur, Rashad Nawfal, Shahabeddin Sotudian, Karl Semaan, Marc Eid, Ji-Heui Seo, Noa Phillips, John J. Canniff, Hunter Savignano, Surya B. Chhetri, Zhenjie Jin, Yukun Ou, Mohamed-Amine Bani, Gwo-Shu Mary Lee, Rachel Trowbridge, Ilana B. Epstein, Gabriella Rickards, Paulo Roberto Da Silva Cordeiro, Ze Zhang, Razane El Hajj Chehade, Brady James, Christophe Massard, Antoine Italiano, Antoine Hollebecque, Jean-Charles Soria, Fabrice André, Cécile Badoual, Joaquim Bellmunt, Harshabad Singh, Andrew J. Aguirre, Brian M. Wolpin, Toni K. Choueiri, Sylvan C. Baca, Matthew L. Freedman

## Abstract

Gene expression is a defining determinant of tumor identity, behavior, and therapeutic response, yet remains challenging to measure noninvasively. Here, we introduce APEX (Associating Plasma Epigenomic features with eXpression), a framework for inferring expression from circulating cell-free chromatin. Trained on ∼270,000 gene–sample pairs from matched tumor RNA-seq and plasma cfChIP-seq across multiple cancers and validated on >15 unseen cancer subtypes, APEX accurately infers cancer gene expression across a range of tumor fractions and outperforms existing plasma-based approaches by integrating positional histone mark and DNA fragmentation patterns across promoters and gene bodies. Using plasma alone, APEX enables classification of prognostically relevant basal and classical pancreatic cancer subtypes and identifies plasma-inferred *NECTIN4* expression as a biomarker of response to enfortumab vedotin in metastatic bladder cancer. Together, these findings establish APEX as a biopsy-free approach for profiling tumor transcriptional states and extend liquid biopsy beyond genomic alterations to clinically relevant gene expression programs.

## MAIN TEXT

Gene expression is the functional output of the DNA code that shapes cellular phenotype and function. In cancer, genetic alterations and their downstream effects on transcription and protein expression provide key insights into tumor biology and therapeutic vulnerabilities^1^. With the rapid development of targeted biologics against variably expressed cell-surface proteins^2^ and increasing evidence for non-genetic mechanisms of therapy resistance^3^, there is a growing need for dynamic and accessible measures of gene expression for cancer patients^4^.

Current approaches rely largely on invasive tissue biopsies, which are limited by sampling bias, spatial heterogeneity, and the inability to longitudinally monitor tumors. Liquid biopsy offers a promising noninvasive alternative, but most cell-free nucleic-acid-based assays focus on mutation detection, with limited tools available to infer transcriptional activity from blood^5^.

Although cell-free RNA (cfRNA) profiling can report on tissue and cell-type composition, existing approaches are constrained by RNA instability, heterogeneous origins, and technical variability^6,7^, motivating complementary strategies that infer gene expression from the more stable chromatin features of cell-free DNA^6^.

Cell-free DNA circulates as protein-protected fragments that reflect the chromatin and regulatory state of their cells of origin^8^. Multiple cfDNA fragment features have been linked to gene expression, including promoter-proximal fragment length entropy^9,10^, fragment length and position distributions^11^, end-motif sequence frequencies^12,13^, enrichment of transcription factor–sized fragments (<80 bp)^14^, and depletion of nucleosome-sized fragments (120–180 bp)^9,12^.

While these DNA-only approaches are simple and cost-effective, they lack molecular enrichment for regulatory or tumor-specific fragments, leading to substantial background signal and often require deep sequencing. Alternatively, cfDNA can be enriched for regulatory signal by immunoprecipitating specific epigenomic marks, including DNA methylation and histone modifications^8^. However, the complementary relationship between immunoprecipitated fragmentomic features and gene regulation has not been explored.

Distinct patterns of histone modifications across the genome are well-established correlates of local gene expression^15^. Recent advances in cell-free chromatin immunoprecipitation sequencing (cfChIP-seq) have enabled genome-wide, noninvasive profiling of histone modifications linked to gene regulation, including H3K4me3 at promoters, H3K27ac and H3K4me2 at promoters and enhancers, H3K9me3 in heterochromatin, and H3K36me3 across actively transcribed gene bodies^16–19^. Existing methods for inferring gene expression from epigenomic data often lack continuous quantitative output, are limited to subsets of genes, provide poor interpretability, or lack external validation (**Supplementary Table 1**). Notably, no approaches are designed to infer gene expression from cfChIP-seq data.

Here, we introduce APEX (Associating Plasma Epigenomics with eXpression), a machine learning framework that integrates histone modification and fragmentomic features from circulating chromatin to infer genome-wide expression noninvasively. Trained on a pan-cancer cohort with matched plasma cfChIP-seq and tumor RNA-seq, APEX leverages promoter- and gene body-associated features to generate accurate and interpretable transcriptional estimates. We validate APEX across multiple independently generated cohorts, demonstrating robust performance across cancers and benchmarking tasks, and with improved accuracy over existing chromatin- and fragmentomic-based approaches. Finally, we show that APEX enables downstream applications including cancer subtyping, differential expression and pathway analysis, and identification of therapeutically actionable targets. Together, these findings position APEX as a broadly applicable approach for noninvasive transcriptional profiling of cancer that can guide treatment selection and monitor dynamic changes in expression as tumors evolve.

## RESULTS

### Benchmarking gene body marks for cell-free chromatin immunoprecipitation

We previously developed a method for immunoprecipitating and sequencing plasma-derived DNA fragments marked by H3K4me3 (promoters) and H3K27ac (promoters and enhancers) from 1 mL of stored plasma^16^. We reasoned that incorporating histone marks spanning distinct genomic regions could further enhance signal detection and resolution beyond these promoter-and enhancer-associated features. Because gene body-associated histone marks occupy nearly tenfold more of the mappable human genome compared to promoter and enhancer marks and are highly predictive of gene expression in human tissues (**Extended Data Fig. 1a,b**), we benchmarked active histone marks enriched within this region. We screened six histone marks in 1 mL of plasma from cancer-free volunteers and patients with prostate cancer, as well as low-input LNCaP chromatin. We identified H3K36me3 and H4K20me1 as the marks most enriched in gene bodies and best able to distinguish prostate cancer from non-cancer chromatin based on signal at differentially expressed genes (**Extended Data Fig. 1c–g; Supplementary Table 2**). Based on its broad genomic occupancy, strong correlation with gene expression, and high gene body and lineage-specific enrichment, we selected H3K36me3 for profiling and included it in downstream analysis.

### In silico screen of cell-free chromatin features linked to cancer gene expression

To systematically evaluate cfChIP-seq features associated with cancer gene expression, we analyzed samples from 15 patients across seven cancer types enrolled in the MOSCATO–1 clinical trial^20^, each with matched plasma cfChIP-seq profiling of H3K4me3, H3K27ac, and H3K36me3 (ichorCNA-estimated tumor fraction^21^, <3% to 42%; median 14%) and tumor RNA-seq collected within 24 hours (**Fig. 1a; Supplementary Table 3**).

**Figure 1.**
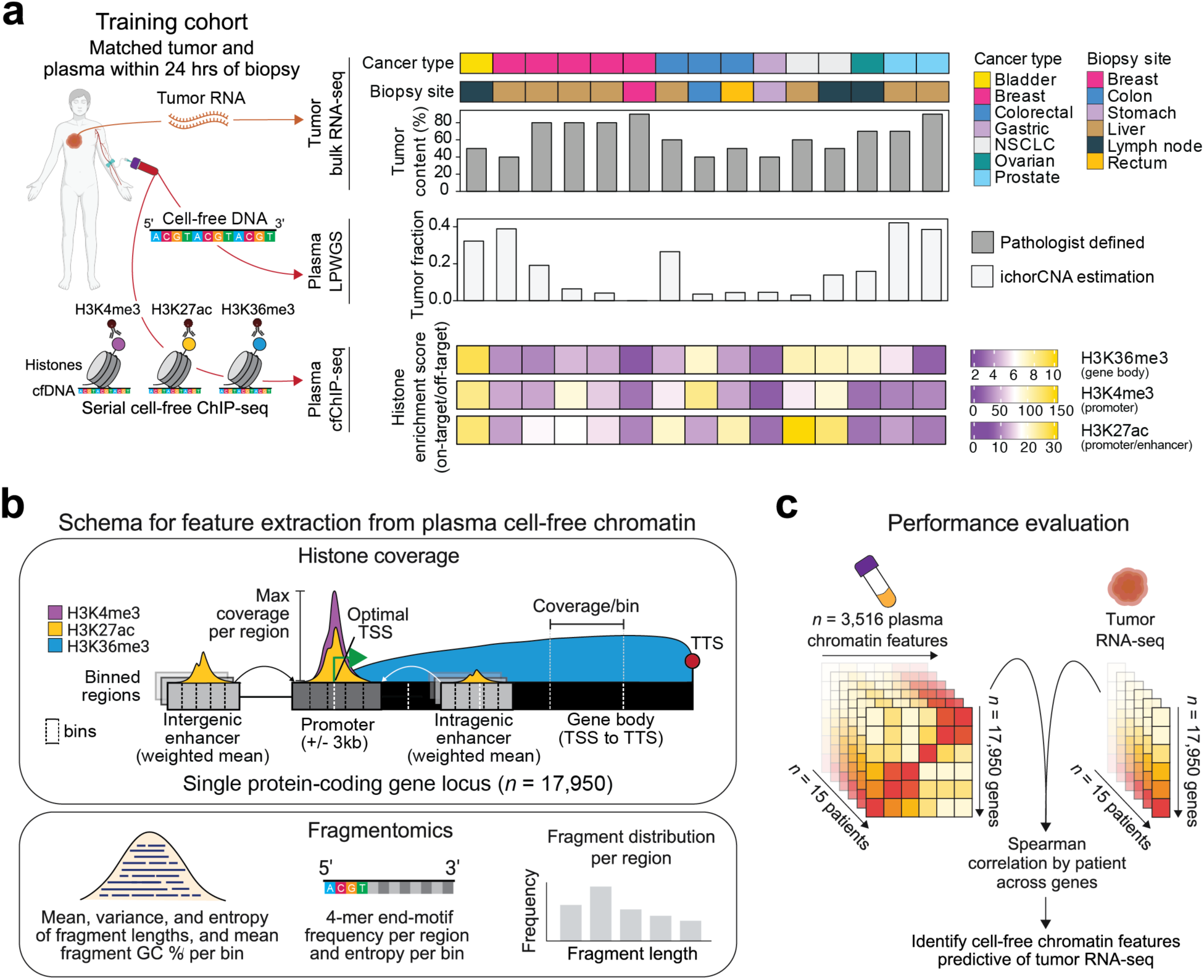
Framework for correlating cell-free chromatin features in plasma with cancer gene expression profiles. **a,** Overview of the training cohort of matched tumor and plasma samples collected within 24 hours across multiple cancer types and biopsy sites. Samples were profiled by bulk tumor RNA sequencing (RNA-seq), plasma low-pass whole-genome sequencing (LPWGS), and serial plasma cell-free ChIP-seq targeting H3K4me3, H3K27ac, and H3K36me3. Sample characteristics, including cancer type, biopsy site, tumor content (%), tumor fraction, and histone enrichment scores are shown (**Supplementary Table 3**). **b,** Schema for extracting plasma cell-free chromatin metrics across 17,950 protein-coding genes. For each gene, 3,516 features were derived, including binned (5 bins per region) and maximum coverage of H3K4me3, H3K27ac, and H3K36me3 across promoter, gene body, intergenic, and intragenic enhancers, as well as fragmentomic measures, including fragment length distribution, 4-mer end motifs, and fragment GC content, summarized by bin or region (**Methods**). The optimal transcription start site (TSS) was selected based on maximum H3K4me3 among candidate TSS (**Methods**). TTS, transcription termination site. **c,** Strategy for evaluating plasma cell-free chromatin metrics most informative of tumor transcriptomic activity. Spearman correlation was used to associate 3,516 plasma chromatin features with matched tumor gene expression (RNA-seq) across 17,950 genes in 15 patients.

We then conducted an in silico screen of chromatin features across ∼18,000 protein-coding genes and their regulatory elements (**Fig. 1b,c**; **Fig. 2a; Supplementary Table 4**). For each gene, we defined promoter (±3 kb around the transcription start site [TSS]), gene body (TSS to transcription termination site [TTS]), and, where annotated, intra- and intergenic enhancer regions based on GeneHancer associations^22^ (**Supplementary Table 5**), each subdivided into equal-sized bins to capture fine-scale variation in epigenomic patterns (**Fig. 1b**). From these regions, we derived coverage- and fragment-based chromatin features and assessed their association with transcriptional activity by computing pairwise Spearman correlations between each plasma cfChIP-seq feature and matched tumor RNA-seq expression across ∼18,000 genes (**Fig. 1c**; **Fig. 2a**).

**Figure 2.**
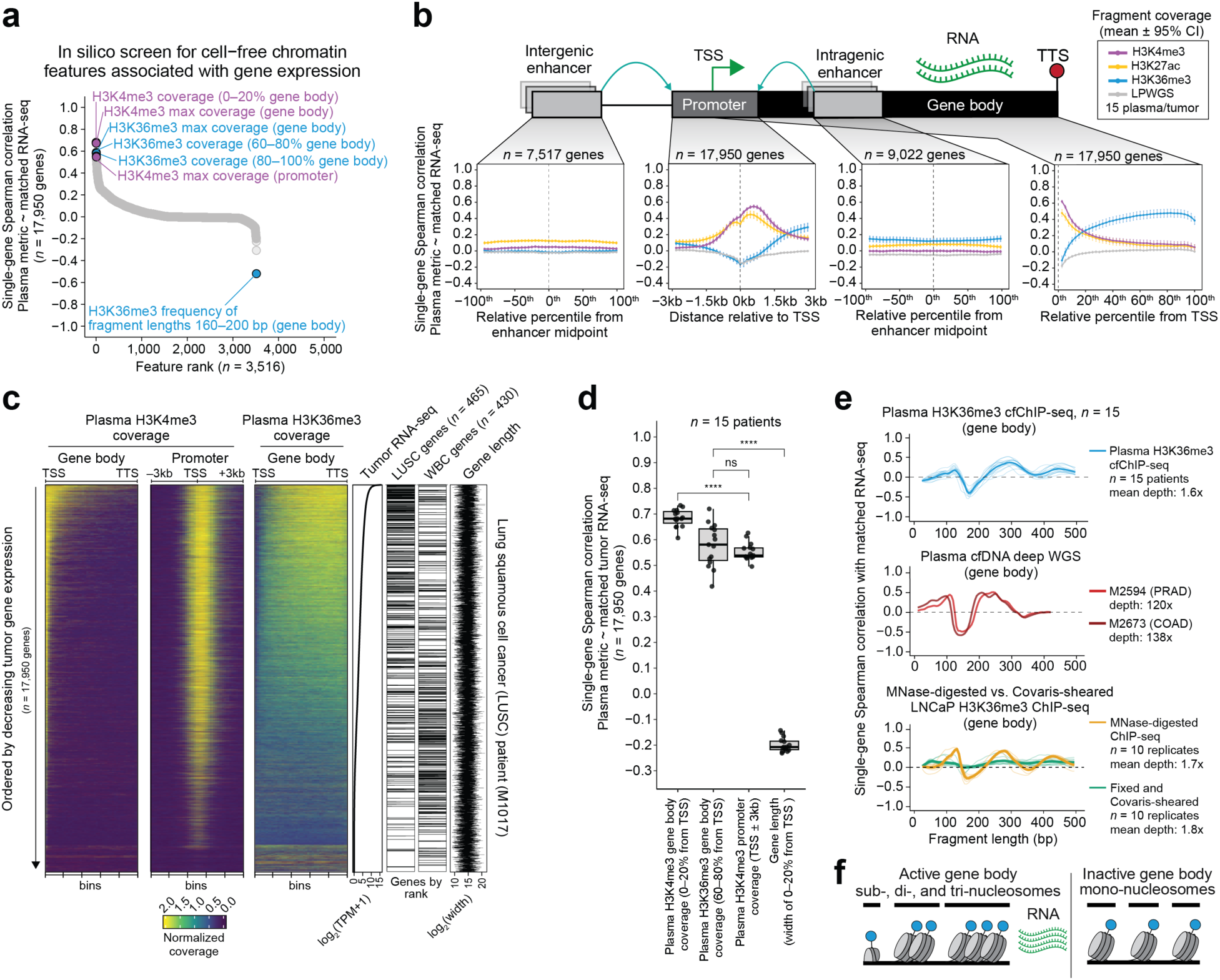
Plasma cell-free chromatin features associated with gene expression across cancer types. **a,** Scatter plot showing results from a systematic screen of plasma cell-free chromatin features from H3K4me3, H3K27ac, and H3K36me3 cfChIP-seq correlated with matched tumor RNA-seq. Each point represents one of 3,516 features, rank-ordered by single-gene Spearman correlation across 17,950 evaluable genes. The most positively and negatively correlated features are highlighted (**Supplementary Table 4**). **b,** Association between plasma histone coverage and matched tumor RNA-seq across genomic regions for 15 patients. Line plots show mean single-gene Spearman correlations (points) with 95% confidence intervals (vertical bars). Plasma-derived histone coverage includes H3K4me3 (purple), H3K27ac (yellow), H3K36me3 (blue), and low-pass whole-genome sequencing (LPWGS; gray). Correlations were computed in 40 bins spanning intergenic enhancers, intragenic enhancers, promoters, and gene bodies, as shown schematically above. For enhancers, coverage was calculated as the confidence-weighted mean across all annotated peaks per gene and bin. The vertical dotted line marks the transcription start site (TSS). Numbers of evaluable genes per region are shown above each box. Enhancer windows were normalized into 40 bins spanning −100 to 100 percentiles around the midpoint; promoter windows were divided into 40 bins across ±3 kb from the TSS; and gene bodies were partitioned into 40 percentile bins from TSS to transcription termination site (TTS). **c,** Heatmaps showing normalized plasma H3K4me3 coverage in the gene body (left) and promoter (middle), and normalized H3K36me3 coverage in the gene body (right) across 17,950 genes, ordered by decreasing tumor RNA-seq signal (line plot) for a representative patient with lung squamous cell carcinoma (LUSC). Rank of LUSC- and white-blood-cell-specific genes and gene length (line plot), in the same gene order, are shown on the far right. **d,** Boxplots showing single-gene Spearman correlations between tumor RNA-seq and plasma cell-free chromatin features across 17,950 evaluable genes in 15 patients. Compared features include H3K4me3 coverage in the promoter (±3 kb from the TSS) and in the first 20% of the gene body; H3K36me3 coverage in the 60–80% gene-body region; and genomic length corresponding to the first 20% of the gene body. Statistical significance was assessed using two-sided paired Wilcoxon tests (\*\*\*\**P* < 0.0001; ns, not significant). **e**, Line plots showing the relationship between fragment length (*x*-axis) and single-gene Spearman correlation with matched tumor RNA-seq (*y*-axis). Top, plasma H3K36me3 cfChIP-seq fragments from 15 patients (individual, light blue; mean, dark blue). Middle, plasma deep whole-genome sequencing (WGS) fragments from two additional patients not in the training cohort (shades of red). Bottom, fragments from LNCaP prostate cancer cells digested with micrococcal nuclease (orange; *n* = 10) or fragmented by Covaris ultrasonication (green; *n* = 10). **f**, Model illustrating how nucleosome organization and fragmentation patterns distinguish transcriptionally active from inactive chromatin regions in cfDNA.

Consistent with prior work^16^, histone coverage showed robust positive correlations with gene expression, whereas low-pass whole genome sequencing, a negative control for copy number-driven effects, exhibited weak or no correlation (**Fig. 2b**). Fragment-length Shannon entropy, 4-mer end-motif diversity, and promoter-associated GC content were modestly correlated with expression (**Extended Data Fig. 2a–c**), while fragment-length mean and variance showed little association (**Extended Data Fig. 2d,e**). No individual 4-mer sequence exhibited associations beyond those attributable to underlying GC content (**Extended Data Fig. 2f**).

Notably, H3K4me3 coverage downstream of the transcription start site (TSS) was more strongly associated with matched tumor gene expression than upstream coverage (**Fig. 2b–d**). Enrichment of H3K4me3 within the proximal gene body (first 0–20% downstream of the TSS), independent of gene length, predicted cancer expression levels, consistent with prior links to lineage-specific and oncogene-associated transcriptional programs^23^ (**Fig. 2c**).

H3K36me3 coverage in the 3′ region of the gene body (60–80% downstream of TSS) also correlated with gene expression (**Fig. 2c,d**), in agreement with prior descriptions^24^. Within H3K36me3-associated fragments in the gene body, enrichment of subnucleosomal (80–120 bp) and di-/tri-nucleosomal (280–320 and 400–440 bp) fragments, and depletion of mononucleosomal (160–200 bp) fragments, were strong correlates of gene expression (**Fig. 2e, *top*; Extended Data Fig. 3**). A similar pattern between fragment lengths and gene expression was observed in deeply sequenced cfDNA (120× and 138×) from two cancer patients with matched tumor RNA-seq (**Fig. 2e, *middle***)^9,11^.

To assess whether these fragment-expression associations reflect nucleosome organization, we analyzed LNCaP cell nuclei subjected to either micrococcal nuclease (MNase) digestion or sonication (**Methods**). MNase digestion selectively cleaves linker DNA between nucleosomes, preserving native chromatin architecture, while sonication randomly shears DNA and disrupts nucleosomal context. In MNase-treated nuclei, fragment-length distributions closely resembled those in plasma and correlated with gene expression from matched RNA-seq, whereas sonicated nuclei lacked this relationship (**Fig. 2e, *bottom*)**. Together, these findings show that fragmentation features of cfDNA enriched by cfChIP-seq associate with transcriptional activity (**Fig. 2f**).

### Development of APEX

Building on the complementary information encoded across promoter, enhancer, and gene body chromatin features, we developed a framework to predict genome-wide gene expression from cell-free epigenomic profiles. We used 3,516 cfChIP-derived positional coverage and fragmentomic features (**Fig. 2**; **Supplementary Table 6**) to train an extreme gradient boosting (XGBoost) regression model to predict matched tumor RNA-seq expression across ∼270,000 gene–sample pairs (∼18,000 genes × 15 patients) (**Fig. 3a; Extended Data Fig. 4a**). XGBoost was selected among multiple regression methods after benchmarking and given its ability to handle multicollinearity among correlated chromatin features (**Extended Data Fig. 4b**). Model training incorporated percentile scaling of input features and cross-validated tuning of region and model parameters to minimize error relative to matched tumor gene expression (log₂[TPM+1]) (**Extended Data Fig. 4c–o**). As many genes use alternative promoters depending on cellular context^25^, we also dynamically reassigned each gene’s TSS per sample based on the position of maximal H3K4me3 signal among annotated TSSs, which significantly improved the performance of promoter-based coverage features compared to using default coordinates (**Extended Data. Fig. 5; Supplementary Note**)

**Figure 3.**
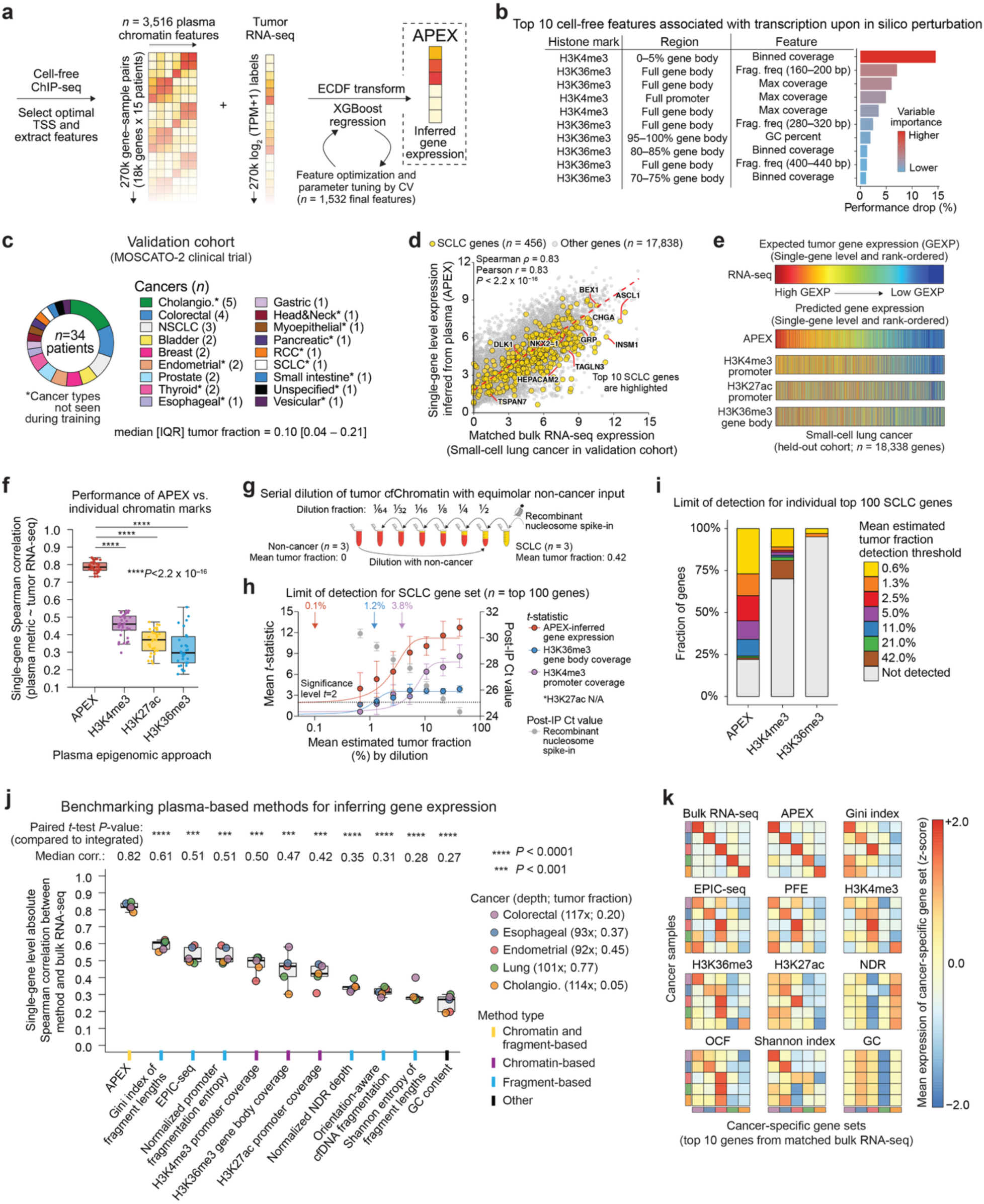
Development and benchmarking of APEX. **a,** Overview of the APEX model (**Methods**). **b,** Top ten plasma cell-free chromatin features ranked by their contribution to model performance, quantified by the relative drop in accuracy upon random permutation. **c,** Summary of the validation cohort of 34 patients encompassing additional tumor types (**Supplementary Table 7**). **d,** Scatter plot showing APEX-predicted (*y*-axis) versus measured tumor RNA-seq (*x*-axis) gene expression across 18,338 genes in a representative small-cell lung cancer (SCLC) sample from the validation cohort. 456 SCLC-specific genes are shown in yellow, with the ten most specific genes labeled. Correlation coefficients and corresponding *P*-values are shown in the top left. The red line denotes the linear regression fit. **e,** Heatmaps comparing expected tumor RNA-seq and APEX-predicted gene expression profiles alongside individual histone coverage features, including H3K4me3 promoter, H3K27ac promoter, and H3K36me3 gene body coverage, at single-gene resolution for the same SCLC sample in **d**. **f,** Box plots showing single-gene Spearman correlations between APEX, plasma histone coverage, and matched tumor RNA-seq across 34 validation samples. Statistical significance was calculated by paired Wilcoxon test between APEX and plasma histone coverage metrics. \*\*\*\**P* < 0.0001. **g–i,** Measuring the limit of detection for cell-free chromatin-based methods. **g,** Schematic of the limit-of-detection experiment in which cfChromatin from three SCLC patients was serially diluted with equimolar cfChromatin from three non-cancer individuals to experimentally mimic different tumor fractions. Recombinant nucleosomes were spiked into SCLC plasma to confirm dilutions. **h,** Detection of a 100-gene SCLC signature, normalized to a WBC-derived background, across serial cfChromatin dilutions relative to non-cancer baseline, as measured by a two-sided *t*-test. For each dilution step, the *x*-axis reflects the mean effective tumor fraction derived from the three starting tumor fractions (0.31, 0.41, 0.55), and the *y*-axis shows the mean *t*-statistic ± SEM across biological replicates. Sigmoid curves were fit to these mean values, and a threshold of *t* = 2 was used to estimate analytical sensitivity. Colors indicate evaluated metrics, including qPCR Ct values for recombinant spike-in nucleosomes (gray). Endogenous H3K27ac signal was insufficient for quantitative analysis. **i,** Detection of 100 individual SCLC genes shown as the stacked bar plot of the lowest significant tumor fraction at which each gene is significantly enriched (*P* < 0.05) above a background distribution of 443 WBC-signature genes**. j,** Benchmarking APEX against alternative plasma-based methods for inferring gene expression using matched plasma and tumor RNA-seq from five patients in the validation cohort. Plasma samples were profiled by cfChIP-seq and deep WGS for fragmentomic analyses. Single-gene Spearman correlation across 18,388 genes served as the benchmark. Legends for samples (colored points) and methods (colored *x*-axis ticks) are shown on the right. Median single-gene Spearman correlations and paired *t*-test *P*-values are shown above. **k,** Heatmaps showing the mean expression of the top ten cancer type-specific genes, defined from matched bulk tumor RNA-seq (top left), as predicted by each plasma-based method. For each cancer type, expression of the corresponding gene signature was *z*-score normalized, and each square represents the normalized expression of that signature in the indicated tumor type. Colored bars and method labels correspond to those in **j**. Box plots in **f** and **j** show medians, quartiles, and 1.5 × IQR. NDR, nucleosome-depleted region; TF, tumor fraction; IP, immunoprecipitation.

To assess the contribution of individual variables to model performance, we systematically ablated regions, histone marks, and feature groups and evaluated prediction accuracy using leave-one-out cross-validation (LOOCV) in the training cohort (**Extended Data Fig. 6**). Both H3K4me3 and H3K36me3 significantly contributed to expression prediction, whereas H3K27ac did not improve performance (**Extended Data Fig. 6a**). Promoter and gene-body features were more informative than enhancer features (**Extended Data Fig. 6b**), and inclusion of DNA methylation provided a statistically significant but quantitatively minor improvement (**Extended Data Fig. 6c**). Removal of individual feature groups demonstrated that no single group recapitulated the performance of the full integrated model (**Extended Data Fig. 6d**). Feature importance analysis by random permutation confirmed H3K4me3 gene body signal (0–5% of the gene body), H3K36me3 fragment length 160–200, and maximum H3K36me3 gene body coverage as top predictors (**Fig. 3b; Supplementary Table 6**).

Based on these analyses, the final model, termed APEX [for Associating Plasma Epigenomics with eXpression] was simplified to 1,532 promoter- and gene-body-derived coverage and fragment-based features from H3K4me3 and H3K36me3 only. This focused model yielded comparable accuracy to the full feature set while substantially improving computational efficiency and assay feasibility (**Extended Data Fig. 6e–h**). APEX achieved a median single-gene Spearman correlation of 0.80 under LOOCV in the training and generalized across tumor types, with performance remaining robust to noise arising from variability in RNA-seq measurements and across a wide range of training set sizes (**Extended Data Fig. 7a–d**).

### Performance of APEX

To validate our findings, we applied APEX to 34 held-out matched plasma and tumor samples, including 18 cancer types with comparable tumor fraction as the training cohort, 11 of which are not represented in the training set (**Fig. 3c; Extended Data Fig. 7e; Supplementary Table 7**). The model consistently demonstrated high correlations with matched tumor RNA-seq across cancer types, achieving comparably high performance compared to the training cohort and outperforming individual histone mark coverage (**Fig. 3d–f; Extended Data Fig. 7f**).

Performance was not restricted to white blood cell (WBC) background, defined as genes expressed at higher levels in whole blood compared to other tissues based on public RNA-seq data, and remained high for non-WBC genes, with correlation performance being superior to individual histone marks across tumor fractions (**Extended Data Fig. 7g,h**).

To directly evaluate the effect of tumor fraction on APEX’s ability to detect cancer-specific signal, we serially diluted tumor-derived plasma cell-free chromatin from three separate patients with small-cell lung cancer with equimolar non-cancer donor plasma (**Methods; Fig. 3g**). We then quantified the limit of detection for SCLC gene expression using a published approach^14^.

Compared to individual histone marks, APEX detected expression of a SCLC signature at a lower tumor fraction (0.1% vs 1.2% and 3.8%; **Fig. 3h; Supplementary Table 8**) and detected more individual SCLC genes, such as *CHGA* and *DLL3*^26,27^ (78% vs 30% and 5%; **Fig. 3i; Extended Data Fig. 8**), at a statistically significant threshold.

To benchmark APEX against existing methods for inferring gene expression, we selected five samples from the validation cohort with sufficient cell-free DNA (20–100 ng) and performed cfChIP-seq and deep whole-genome sequencing (mean: 103× coverage) (**Supplementary Table 9**). APEX-inferred gene expression significantly outperformed existing cfChIP-seq and fragmentomic approaches in recovering gene expression (median Spearman correlation with matched tumor RNA-seq vs second-best performer^9^: 0.82 vs. 0.61; *P*<0.0001), including improved discrimination between cancer-specific expression programs (**Fig. 3j,k**). While deeper sequencing may improve the performance of other methods, these results demonstrated that combining chromatin and fragmentomic features using APEX substantially outperforms any individual feature.

### Applications of APEX

Accurate cancer subtyping is critical for diagnosis, prognosis, and treatment selection. To assess APEX’s ability to distinguish cancer subtypes, we applied it to 132 previously published H3K4me3 profiles spanning eight cancer subtypes (**Supplementary Table 10**)^16^. As H3K36me3 data were not available for this dataset, we used APEX with H3K4me3 alone. We evaluated subtype-specific gene expression using Wilcoxon tests against RNA-seq signatures derived from public data matched for tumor subtype (**Supplementary Table 8**). Even with a single histone mark, APEX–H3K4me3 significantly outperformed promoter-aggregated H3K4me3 signal, underscoring its utility in distinguishing tumor subtypes based on gene expression differences (**Fig. 4a,b**).

**Figure 4.**
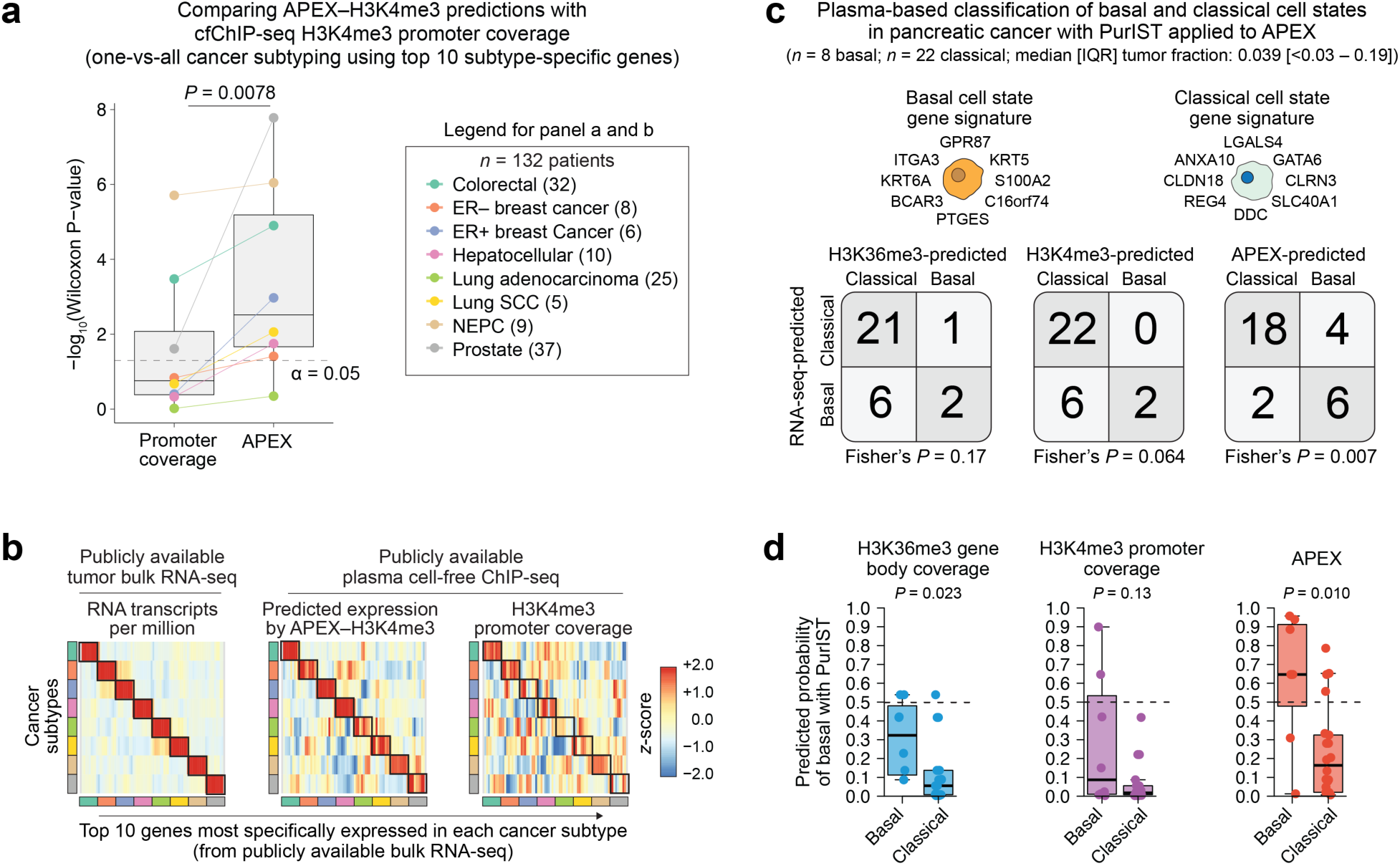
APEX discriminates cancer subtypes and transcriptional cell states from plasma. **a,** Comparison of APEX using only H3K4me3 input (APEX–H3K4me3) with plasma cfChIP-seq H3K4me3 promoter coverage in a one-vs-all cancer subtype classification task across eight subtypes from 132 previously profiled patients^16^. The plot shows –log(Wilcoxon *P*-values) for discrimination accuracy using the top ten subtype-specific genes defined from subtype-matched publicly available bulk RNA-seq (subtypes listed at right). The horizontal gray dashed line indicates the α = 0.05 significance threshold. ER, estrogen receptor; SCC, squamous cell carcinoma; NEPC, neuroendocrine prostate cancer. **b,** Heatmaps showing signal from the top ten genes for each cancer subtype across three data layers: tumor RNA-seq expression (transcripts per million; left), APEX-inferred H3K4me3 expression from plasma (center), and plasma cfChIP-seq H3K4me3 promoter coverage (right). Colored bars indicate subtypes as shown in panel **a**. For visualization only, values were *z*-score normalized and smoothed using a centered rolling mean (window size = 3) along the cancer-subtype axis. **c**, Plasma-based classification of basal (*n* = 8) and classical (*n* = 22) pancreatic cancer cell states using PurIST^28^ applied to H3K4me3 promoter coverage, H3K36me3 gene-body coverage, or APEX predictions, compared against matched tumor RNA-seq ground truth. Basal and classical PurIST gene signatures are shown schematically. 2 × 2 contingency tables with Fisher’s exact test *P*-values are displayed. **d,** Box plots showing the predicted probability of the basal cell state from the PurIST logistic regression model as in **c**. The horizontal dashed line at 0.5 indicates equal probability of basal versus classical states. Box plots in panels **a** and **d** show medians, quartiles, and 1.5 × IQR.

Because APEX outputs inferred gene expression, it enables the direct application of established RNA-seq-based analytical frameworks to plasma data. PurIST is a validated single-sample transcriptional classifier that assigns pancreatic tumors to prognostically distinct basal or classical subtypes based on the relative expression of a 16-gene signature, with important implications for patient outcomes and therapeutic sensitivity^28^. We applied PurIST to APEX-inferred gene expression, or to signal from individual histone marks, using plasma from 30 pancreatic cancer patients with tumor RNA-seq-confirmed basal or classical subtypes (**Supplementary Table 11**). APEX enabled significantly more accurate subtype classification than individual histone marks, demonstrating that clinically relevant transcriptional subtypes can be measured noninvasively from plasma-inferred expression profiles (balanced accuracy: APEX = 0.78; H3K36me3 = 0.60; H3K4me3 = 0.63; **Fig. 4c,d**).

### Plasma-inferred *NECTIN4* expression predicts response and survival to enfortumab vedotin in metastatic bladder cancer

Emerging tissue-based studies have shown that tumor *NECTIN4* expression correlates with clinical response to enfortumab vedotin (EV), an antibody-drug conjugate targeting *NECTIN4* in metastatic urothelial carcinoma^29–31^. We therefore hypothesized that *NECTIN4* expression inferred from baseline plasma could noninvasively predict clinical outcomes in patients with metastatic bladder cancer treated with EV monotherapy. To test this, we performed cfChIP-seq on baseline plasma samples and inferred cancer gene expression using APEX (**Fig. 5a,b**; **Supplementary Table 12**).

**Figure 5.**
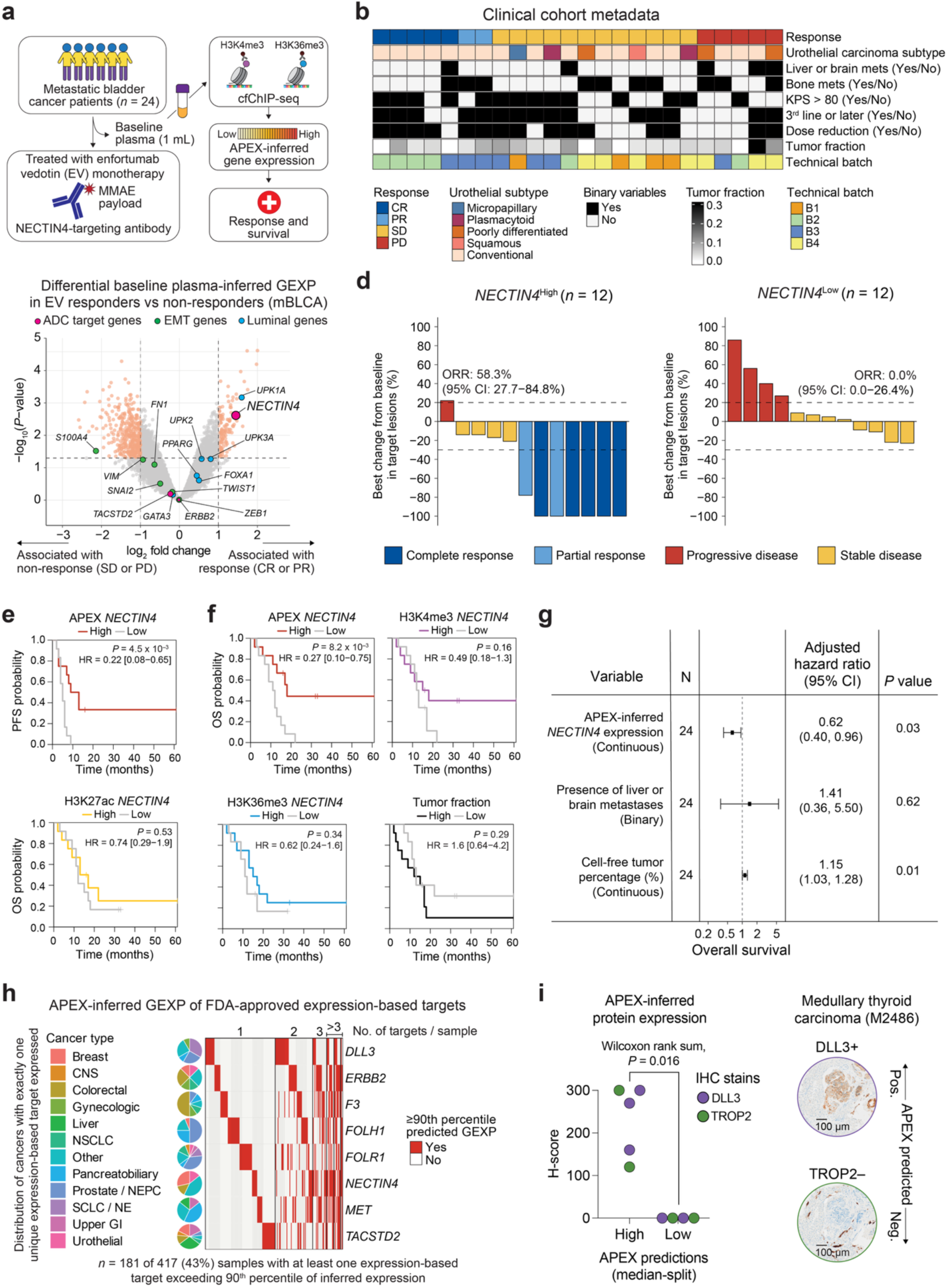
Plasma epigenomic inference of target gene expression predicts antibody-drug conjugate response and survival. **a,** Study overview. Baseline plasma was collected from patients with metastatic bladder cancer prior to treatment with enfortumab vedotin (EV) monotherapy. Plasma cfChIP-seq profiles for H3K4me3 and H3K36me3 were integrated using APEX to infer cancer gene expression and evaluate associations with treatment response and survival. **b,** Clinical and pathological characteristics of the EV-treated cohort (*n* = 24). **c,** Differential baseline APEX-inferred gene expression between responders (complete or partial response) and non-responders (stable or progressive disease). Highlighted are antibody drug conjugate (ADC) targets, epithelial–mesenchymal transition (EMT)-associated genes, and luminal lineage genes. **d,** Best percentage change from baseline in target lesions stratified by high versus low baseline APEX-inferred *NECTIN4* expression (dichotomized by median). Objective response rates (ORR) and 95% confidence intervals are shown. **e, f,** Progression-free survival (**e**) and overall survival (**f**) stratified by high versus low baseline plasma-inferred features, including APEX-inferred *NECTIN4* expression, histone mark-specific cfChIP-seq coverage at the *NECTIN4* locus (H3K4me3 promoter, H3K27ac promoter, H3K36me3 gene body), and plasma tumor fraction estimated using ichorCNA (v0.2.0)^21^. Survival curves were compared using two-sided log-rank tests; hazard ratios (HR) and 95% confidence intervals are shown. **g,** Multivariable Cox proportional hazards analysis for overall survival including continuous APEX-inferred *NECTIN4* expression, presence of liver or brain metastases, and plasma tumor fraction, demonstrating an independent association between inferred *NECTIN4* expression and survival. **h,** Heatmap (right) showing APEX-inferred gene expression (GEXP) of FDA-approved expression-based targets for solid cancers across 417 plasma samples. Red indicates samples with predicted expression at or above the 90th percentile for a given target. Samples (columns) are ordered by the number of cancer targets exceeding the 90th percentile of inferred expression (top). Pie charts indicate the distribution of cancer types among samples with exactly one cancer target exceeding the 90th percentile. **i,** Validation of plasma-inferred protein expression by immunohistochemistry (IHC). *Left*, DLL3 and TROP2 H-scores from tumors profiled in the MOSCATO–2 clinical trial (related to Fig. 3c), stratified by high versus low APEX-inferred expression based on a median split of plasma-inferred values. *Right*, representative IHC images from medullary thyroid carcinoma illustrating concordance between APEX predictions and tissue protein expression. Scale bar, 100 μm.

APEX-inferred *NECTIN4* expression was significantly higher in responders (complete or partial response) compared to non-responders (stable or progressive disease) (*P* = 1.9 × 10^−3^; **Fig. 5c; Extended Data Fig. 9a–d; Supplementary Table 13**). While 58% of patients with plasma-inferred *NECTIN4* expression above the cohort median objectively responded to EV, none with predicted expression below the median did, highlighting the negative predictive value of the assay (**Fig. 5d**).

In contrast, *NECTIN4* estimates derived from individual histone marks showed weaker or nonsignificant associations with response. Plasma-inferred expression of other antibody-drug conjugate (ADC) targets not directly targeted by EV, including *ERBB2* and *TACSTD2*, tumor fraction, and *NECTIN4* copy number were not associated with treatment response, supporting the specificity of the *NECTIN4* signal (**Fig. 5c**; **Extended Data Fig. 9e–j**). Consistent with this finding, responders were enriched for urothelial luminal genes known to associate with *NECTIN4* expression and favorable outcomes^32^, while non-responders showed increased activation of epithelial-mesenchymal transition (EMT)-related genes that have been associated with worse outcomes^33^ (**Fig. 5c**; **Extended Data Fig. 9i; Supplementary Table 8**).

Baseline APEX-inferred *NECTIN4* expression was also significantly associated with both progression-free survival and overall survival. This association was stronger than that observed using individual histone mark coverage, plasma tumor fraction, or *NECTIN4* copy number variation (**Fig. 5e,f**; **Extended Data Fig. 9k,l**). Multivariable Cox proportional hazards models confirmed that APEX-inferred *NECTIN4* remained independently associated with survival after accounting for tumor fraction and metastatic disease burden, supporting its role as an expression-based biomarker for EV sensitivity (**Fig. 5g**).

### Plasma-inferred expression enables nomination of expression-based targets across cancer types

Targeted biologics, including ADCs, multispecific antibodies, radioligand therapies (RLTs), and chimeric antigen receptor (CAR) T cells, are an increasingly important class of cancer therapies guided by cell-surface protein expression rather than recurrent genetic alterations^2^. However, scalable blood-based assays for assessing target expression are limited, particularly in advanced disease where repeated tissue biopsies are impractical despite sufficient circulating tumor DNA.

Leveraging a pan-cancer atlas of cfChIP-seq data (*n* = 417 patients) spanning a range of tumor fractions, we asked whether APEX could identify cell surface targets with exceptionally high expression in a given cancer. Applying APEX to cfChIP-seq data for H3K4me3 alone or combined H3K4me3 and H3K36me3 where available, we quantified the prevalence of high inferred expression across clinically actionable targets. Using a conservative threshold of expression exceeding the 90th percentile across samples, 43% of patients exhibited at least one FDA-approved expression-based target, including *DLL3*, *ERBB2*, *F3*, *FOLR1*, *FOLH1*, *NECTIN4*, *MET*, or *TACSTD2* (**Fig. 5h**).

Consistent with these pan-cancer predictions, plasma-inferred expression of selected targets was supported by tumor immunohistochemistry in the MOSCATO–2 clinical trial, including a representative medullary thyroid carcinoma case with concordant high plasma-inferred DLL3 and low plasma-inferred TROP2 expression relative to tumor protein staining. (**Fig. 5i**). When expanded to include FDA-approved and cell surface targets in clinical development (*n* = 291)^34^ (**Extended Data Fig. 10; Supplementary Table 10**), 97% of patients met this criterion. These data demonstrate how genome-wide assessment of expression from plasma could identify unexpected therapeutic opportunities across diverse cancer types.

## DISCUSSION

In this study, we present APEX, a machine learning framework for noninvasive inference of cancer gene expression from plasma that integrates chromatin and fragmentomic features from cfChIP-seq. By jointly modeling these features, APEX captures complementary aspects of transcriptional regulation that are inaccessible to either modality alone. Further, by leveraging more than 270,000 paired observations of gene expression and local chromatin features across the genome, APEX learns gene-agnostic relationships between epigenomic state and transcription, enabling robust inference in independent cohorts. Systematic screening of plasma-derived chromatin features revealed that proximal gene-body H3K4me3 signal is more informative of expression levels than aggregate promoter-proximal signal, consistent with prior work linking extended H3K4me3 occupancy to lineage-specific and oncogene-associated transcriptional programs^23^. In addition, H3K36me3 signal, particularly in the 3′ gene body and regions of maximal coverage, provided complementary predictive power, reflecting its association with transcriptional elongation^24^.

Beyond histone mark enrichment, APEX leverages fragmentomic features that reflect local chromatin structure. Notably, depletion of nucleosome-sized H3K36me3-immunoprecipitated fragments and relative enrichment of subnucleosomal and di-/trinucleosomal fragments were correlated with gene expression. These observations align with prior studies linking fragment length distributions to cancer-specific chromatin accessibility and mutation burden^9,35^, and suggest that fragment-length features can enhance inference of tumor-specific transcriptional activity beyond coverage-based measures alone.

In contrast, H3K27ac and DNA methylation contributed little to gene-level expression prediction, even when enhancer–gene relationships and strict quality thresholds were applied^22^. This likely reflects a combination of redundancy with promoter- and gene-body-associated marks, greater signal degradation in plasma, and limitations of current enhancer-promoter annotations, which assume homogeneous cell populations and may not capture tissue- and gene-specific enhancer usage in heterogeneous cell-free chromatin mixtures. More precise context-specific enhancer-promoter modeling may improve gene expression prediction, although H3K27ac and DNA methylation remain well established for tumor subtyping and regulatory state classification^16^.

A key advantage of APEX is that it outputs inferred gene expression in a format directly compatible with standard RNA-seq analytical frameworks. This enables extension of established transcriptomic analyses, including differential expression, gene set enrichment, and expression-based classifiers, like PurIST^28^, to liquid biopsy data. Accordingly, we demonstrate improved cancer subtyping and more accurate recovery of lineage-specific transcriptional programs relative to individual histone marks. Although APEX was trained and validated in plasma, it leverages chromatin-expression relationships that are expected to be preserved in other biofluids containing cell-free chromatin^36^.

Our study has several limitations. Although APEX improves expression inference at lower tumor fractions relative to individual histone features, signal acquisition remains unstable at very low tumor fractions (<0.5%). Accordingly, our method is not intended for minimal residual disease detection or early cancer screening, but is most applicable in metastatic disease at diagnosis or progression, where median tumor fractions are closer to ∼3%^37^. While expression predictions retain some dependence on tumor fraction, explicit linear adjustment risks removing biologically meaningful signal, and APEX predictions were empirically stable without tumor-fraction correction. As with other cfChIP-seq-based approaches, pre-analytical and technical variables, including sample handling, freeze–thaw cycles, and antibody performance, may influence data quality, although robust performance across multiple, diverse cohorts suggests resilience to such variation. In addition, APEX was trained using matched single-site RNA-seq as a reference, which may incompletely capture heterogeneity across metastatic lesions; nevertheless, robust performance under label perturbation suggests that APEX captures conserved transcriptional programs encoded in circulating tumor-derived chromatin.

Conceptually, APEX infers transcriptional activity from epigenomic features that reflect active gene regulation rather than direct protein measurements. For many therapeutic targets, transcriptional activity is a major determinant of protein expression, and concordance between plasma-inferred expression and tumor protein abundance was observed for selected targets by immunohistochemistry. However, this validation was limited in scope, and further evaluation on a target-by-target basis will be required to define the relationship between plasma-inferred transcription and tumor protein abundance. While applied to protein-coding genes, APEX can be extended to non-coding transcripts, although this will require further validation.

Clinically, efforts to link cell-surface-targeting therapies, such as ADCs, multispecific antibodies, RLTs, and CAR-T cells, to treatment response have been hindered by reliance on tissue biopsies, which are often archival, spatially heterogeneous, and subject to inter-observer variability in immunohistochemical assessment^4^. This motivated evaluation of APEX as a noninvasive approach for target measurement and nomination. In metastatic bladder cancer, APEX-inferred *NECTIN4* expression predicted response and survival following EV, outperforming tumor fraction, copy-number variation, and single histone mark coverage.

Although tissue-based studies have variably linked tumor NECTIN4 expression to clinical outcomes^30,31,38^, interpretation has been confounded in EV–pembrolizumab regimens^38^ and further limited by heterogeneity across metastatic sites, dynamic changes in expression over time, and the absence of standardized scoring methods or consensus antibodies for assessment^30^. By focusing on EV monotherapy, integrating signals across sites, and measuring expression at treatment initiation, plasma-inferred expression showed stronger associations with outcome. APEX also recapitulated known biological programs associated with favorable and unfavorable outcomes^29,32,33^ and demonstrated specificity for *NECTIN4* relative to other ADC targets. Extending this framework to a large pan-cancer plasma cohort revealed widespread expression of clinically actionable targets, including high-expression outliers in cancer types where corresponding therapies are not routinely used. These findings suggest that plasma-inferred expression can guide rational tissue-based confirmation or inform expression-directed therapeutic decision-making when biopsy is infeasible.

As the repertoire of expression-targeted therapies continues to expand^2^, scalable and noninvasive tools for assessing cancer gene expression will become increasingly important. APEX provides a generalizable framework for plasma-based transcriptional profiling, enabling longitudinal monitoring, expanded access to biomarker evaluation, and systematic nomination of therapeutic targets. Collectively, these results establish APEX as a practical and biologically grounded approach for advancing precision oncology through liquid biopsy-based gene expression inference.

## Acknowledgments

We thank members of the Freedman and Baca laboratories for insightful discussions and feedback throughout the course of this work. We are especially grateful to the patients at Dana-Farber Cancer Institute, whose experiences continue to inspire our research. This work was supported by the Cutler Family Fund for Prevention and Early Detection (M.L.F.), Damon Runyon Cancer Research Foundation (S.C.B.), Dana-Farber/Harvard Cancer Center Kidney SPORE (T.K.C., 2P50CA101942-16) and Program (T.K.C., 5P30CA006516-56), Donahue Family Fund (M.L.F.), Frank Shaughnessy Kidney Cancer Research Fund (T.K.C.), Fund for Innovation in Cancer Informatics (S.C.B.), Hinda and Arthur Marcus Fund (T.K.C.), H.L. Snyder Medical Research Foundation (M.L.F.), Institut Servier (D.V.), Kohlberg Chair at Harvard Medical School and the Trust Family (T.K.C.), Loker Pinard Funds for Kidney Cancer Research at DFCI (T.K.C.), Michael Brigham (T.K.C.), National Cancer Institute (G.S.G, T32CA009172; S.C.B., U01 CA296432; M.L.F., R01 CA262577, R01 CA251555, R01 CA300092, R01 CA259058, P50 CA272390), Pan Mass Challenge (T.K.C.), Philippe Foundation (D.V.), Prostate Cancer Research Foundation (S.C.B.), Roger and Kathy Marino Fund for Research Acceleration (T.K.C.), US Department of Defense (S.C.B., W81XWH-21-1-0358), and Wong Family Translational Oncology Award (S.C.B.).

## Author contributions

G.S.G. conceived the study, designed experiments with guidance from S.C.B. and M.L.F., developed the APEX framework, performed all computational analyses, generated figures, interpreted results, and wrote the manuscript. D.V. led cfChIP-seq experimental efforts, obtained and processed MOSCATO–1 and MOSCATO–2 clinical trial samples, benchmarked gene-body histone marks, and contributed to limit-of-detection analyses. R.N. obtained and processed the enfortumab vedotin cohort, designed and performed associated experiments, curated clinical data, and designed and conducted limit-of-detection studies. S.S. contributed to early conceptual discussions, model design, and statistical analyses. K.S., M.E., H.Sin., A.J.A., and B.M.W. provided pancreatic cancer datasets and contributed to their interpretation. J.-H.S., N.P., J.J.C., and H.Sav. contributed to gene body benchmarking analyses and cfChIP-seq experiments. S.B.C., Z.J., P.R.D.S.C., Z.Z., R.E.H.C., and Y.O. contributed to computational pipeline development. B.J. assisted with cfChIP-seq and DNA processing. G.-S.M.L., J.B., R.T., I.B.E., and G.R. contributed to acquisition and annotation of the EV clinical cohort. C.M., A.I., A.H., J.-C.S., and F.A. contributed MOSCATO samples and matched clinical data. C.B. provided pathology expertise and performed immunohistochemistry. T.K.C., H.Sin., A.J.A., and B.M.W. provided funding and clinical guidance. S.C.B. and M.L.F. supervised the study, provided funding and scientific direction, and edited the manuscript. All authors reviewed and approved the final manuscript.

## SUPPLEMENTARY DATA

Supplementary Data 1 APEX-inferred gene expression profiles for the MOSCATO–1 training cohort.

Supplementary Data 2 Bulk tumor RNA-seq gene expression profiles (logTPM) for the MOSCATO–1 training cohort.

Supplementary Data 3 APEX-inferred gene expression profiles for the MOSCATO–2 test cohort.

Supplementary Data 4 Bulk tumor RNA-seq gene expression profiles (logTPM) for the MOSCATO–2 test cohort.

Supplementary Data 5 APEX-inferred gene expression profiles for limit-of-detection dilution series experiments.

Supplementary Data 6 APEX-inferred gene expression profiles for the metastatic pancreatic cancer cohort.

Supplementary Data 7 Bulk tumor RNA-seq gene expression profiles (logTPM) for the metastatic pancreatic cancer cohort.

Supplementary Data 8 APEX-inferred gene expression profiles for the cohort of metastatic bladder cancer patients treated with enfortumab vedotin.

Supplementary Data 9 APEX-inferred gene expression profiles for samples from a previously published pan-cancer plasma epigenomics study.

---

**Extended Data Figure 1.**
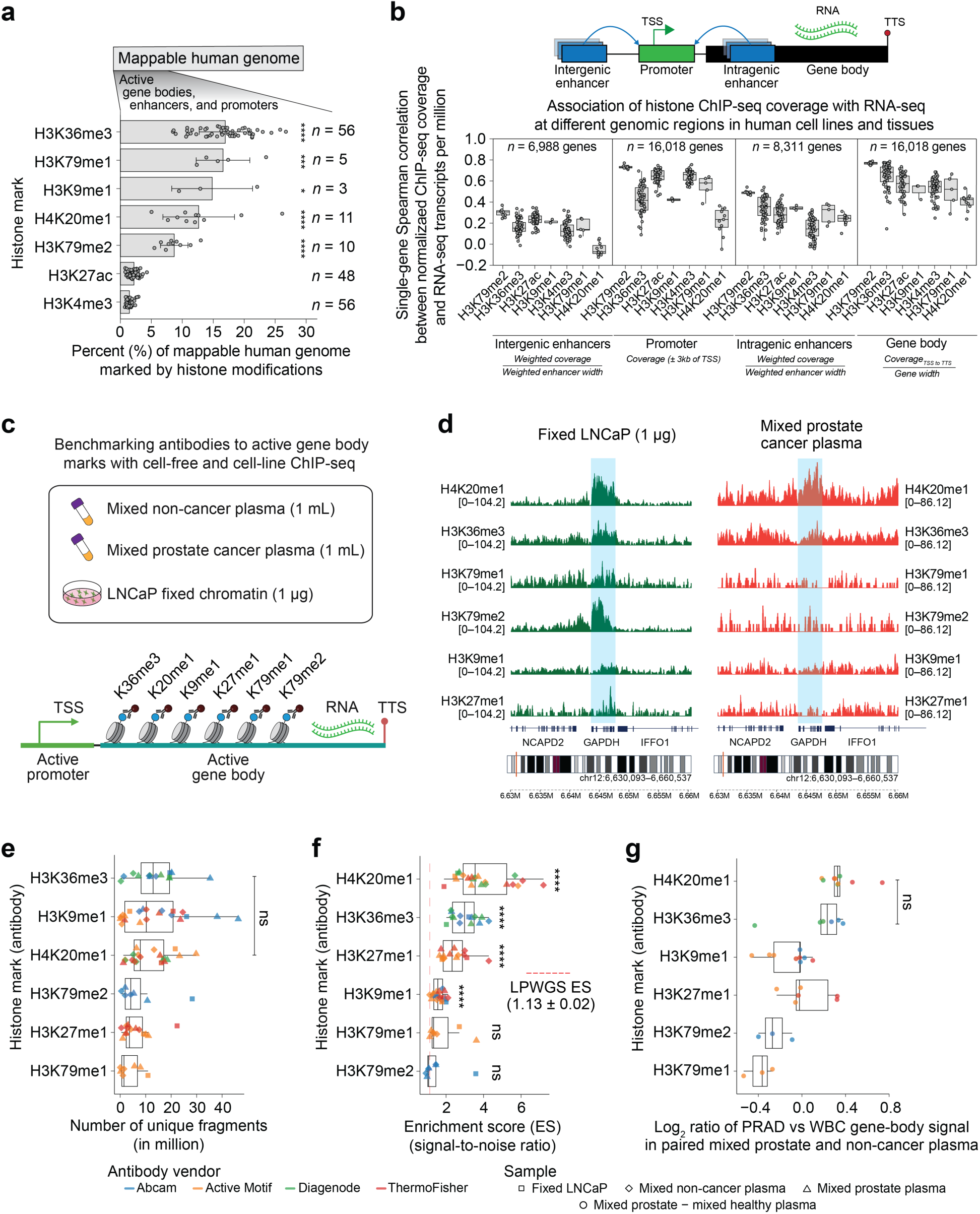
Selection and benchmarking of antibodies to active gene body histone marks in plasma and cell-line ChIP-seq. **a,** Barplots showing mean (solid bars) ± standard deviation (error bars) of the fraction of the genome covered by significant peaks for seven active histone marks across multiple tissues using data obtained from Roadmap Epigenomic tissue ChIP-seq (sample numbers indicated at right). Peaks were called with MACS v2.0.10 using matched whole-cell extract controls, applying narrow-peak thresholds (H3K4me3, H3K27ac; Poisson *P* = 0.01) or broad-peak thresholds (H3K36me3, H3K79me1, H3K79me2, H3K9me1, H4K20me1; Poisson *P* = 0.1) tailored to each mark. Statistical significance between each gene body mark and the promoter-associated mark H3K4me3 was evaluated using an unpaired one-sided *t*-test. **b,** Boxplots showing Spearman correlations between ChIP-seq coverage of seven active histone marks at distinct regulatory regions (intergenic enhancers, intragenic enhancers, promoters, and gene bodies) and RNA-seq transcript abundance. ChIP-seq fragments (tagAlign files) and RNA-seq matrices were obtained from the Roadmap Epigenomics database (sample numbers as in **a**). For enhancers, coverage was calculated as the confidence-weighted mean across all GeneHancer-associated enhancers per gene. Schematic (top) indicates genomic regions relative to gene structure; the number of evaluable genes per region is shown above each box. **c,** Schematic of experimental design for benchmarking antibodies to active gene body marks using cell-free (1 mL of mixed non-cancer and prostate cancer plasma) and low-input (1 µg) cell-line (LNCaP) ChIP-seq. **d**, Signal tracks of ChIP-seq coverage for active gene body marks across an active housekeeping gene, *GAPDH*, in 0.8–1 µg fixed LNCaP chromatin and 1 mL mixed prostate cancer plasma. Individual tracks show different modifications with *y*-axis coverage range indicated in brackets. **e–g,** Performance of ChIP-seq antibodies in fixed LNCaP chromatin and mixed plasma samples, grouped by vendor (color) and sample type (shape). **e,** Number of unique sequencing fragments (millions). **f,** Enrichment score (signal-to-noise ratio), defined as coverage at gene-body regions divided by signal in regions >10 kb from gene bodies; dashed red line indicates threshold from low-pass whole-genome sequencing (LPWGS; non-immunoprecipitated negative control; *n* = 15). **g,** Log_2_ ratio of average gene-body signal across the top 500 prostate (PRAD) versus top 500 white blood cell (WBC) genes in paired mixed prostate cancer and non-cancer plasma. Antibodies are ordered top-to-bottom by median performance. For **e** and **g**, statistical significance was determined with a two-sided unpaired *t*-test between histone marks, while for **f**, statistical significance was determined with a two-sided unpaired *t*-test between histone marks and LPWGS (*n* = 24 cancer plasma samples); \*\*\*\**P* < 0.0001, ns, not significant.

**Extended Data Figure 2.**
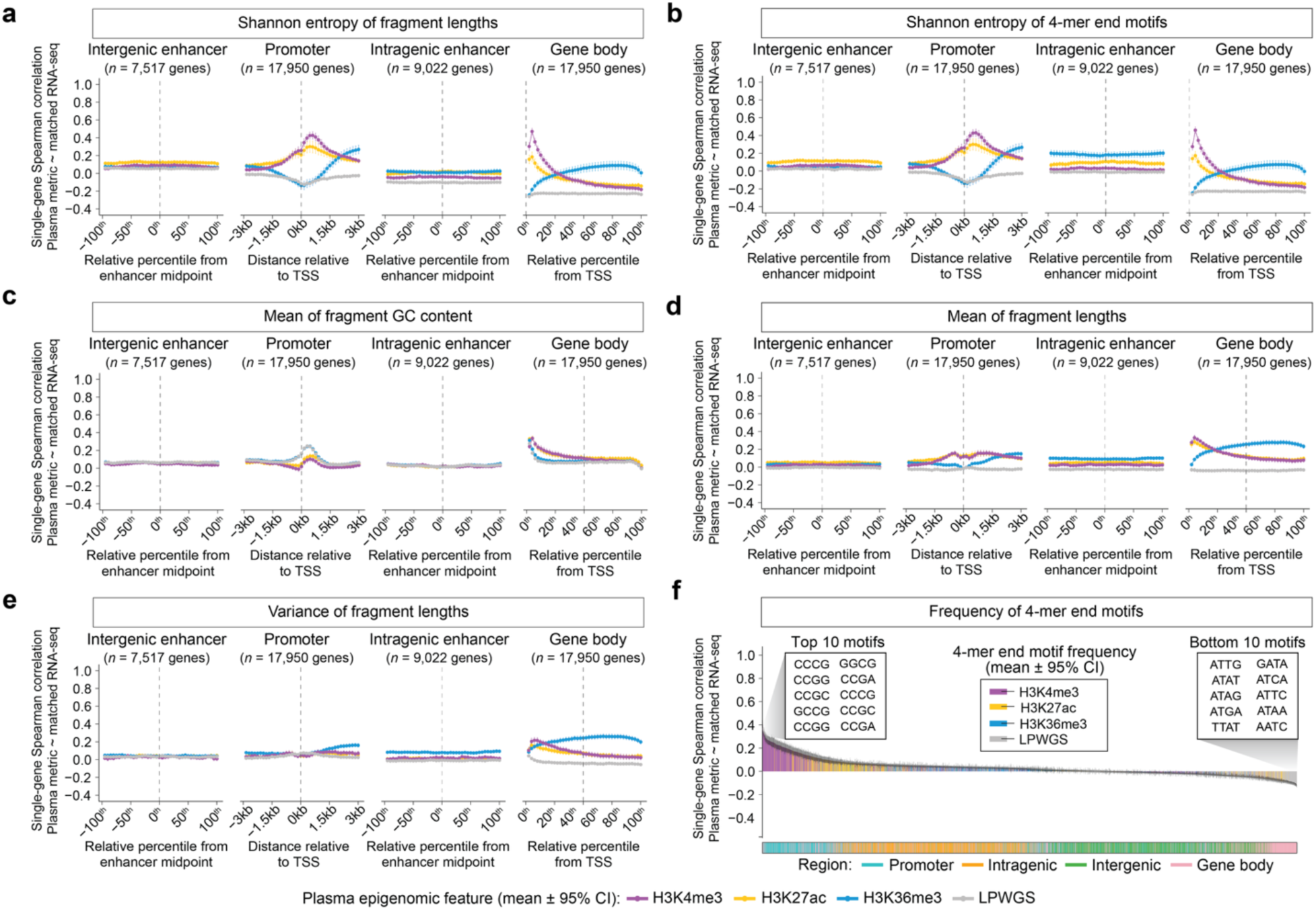
Association between plasma epigenomic features and matched tumor RNA-seq across regulatory regions. **a–e,** Line plots show mean single-gene Spearman correlations (points) with 95% confidence intervals (vertical bars) between plasma-derived epigenomic features and matched tumor RNA-seq across protein-coding genes in 15 patients. Plasma-derived epigenomic features include **a,** Shannon entropy of fragment lengths; **b,** Shannon entropy of 4-mer end motifs; **c,** mean fragment GC content; **d,** mean of fragment lengths; and **e,** variance of fragment lengths. These features were derived from H3K4me3 (purple), H3K27ac (yellow), and H3K36me3 (blue) cfChIP-seq and low-pass whole-genome sequencing (LPWGS; gray). Correlations were computed in 40 bins spanning intergenic enhancers, intragenic enhancers, promoters, and gene bodies. For enhancers, coverage was calculated as the confidence-weighted mean across all annotated enhancer peaks per gene. The vertical dotted line marks the transcription start site (TSS). Enhancer windows were normalized into 40 bins spanning −100 to 100 percentiles around the midpoint; promoter windows were divided into 40 bins across ±3 kb from the transcription start site (TSS); and gene bodies were partitioned into 40 percentile bins from TSS to transcription termination site (TTS). **f,** Barplots showing mean ± 95% confidence intervals for single-gene Spearman correlations between the frequency of each 4-mer end motif and matched tumor RNA-seq, ordered from most positively to most negatively correlated. Frequencies were calculated across promoter, intragenic enhancer, intergenic enhancer, and gene body regions (color-coded at bottom) for 15 patients in the training cohort. The ten 4-mer motifs most positively and negatively correlated with tumor RNA-seq are highlighted.

**Extended Data Figure 3.**
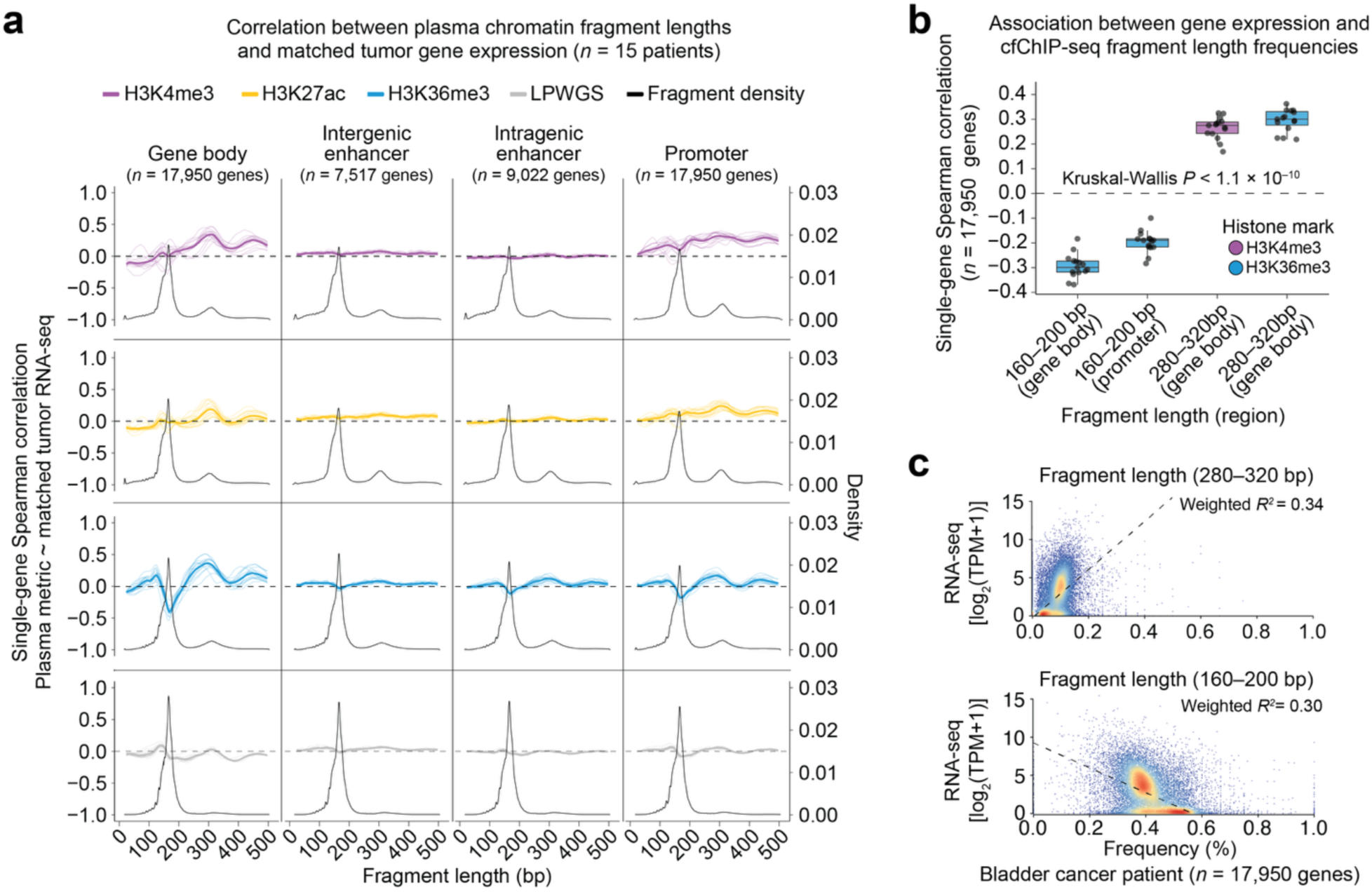
Association between plasma cell-free chromatin fragment length and matched tumor RNA-seq. **a,** Line plots showing the relationship between fragment length (*x*-axis) and single-gene Spearman correlation with matched tumor RNA-seq (*y*-axis) for H3K4me3 (purple), H3K27ac (yellow), H3K36me3 (blue), and low-pass whole genome sequencing (LPWGs; gray) across promoters (transcriptional start site [TSS] ±3 kb), intergenic enhancers, intragenic enhancers, and gene bodies (TSS to transcriptional terminal site [TTS]). Mean correlations across 15 individual samples are shown (dark lines), with per-sample correlations shown in lighter shades. Black lines indicate normalized fragment density profiles, and the horizontal dashed gray line denotes zero Spearman correlation. **b,** Boxplots showing single-gene Spearman correlations between matched tumor RNA-seq and the two most positively and negatively correlated fragment length-related features identified in the in silico screen (corresponding to Fig. 2a; *n* = 15 patients). Statistical significance was assessed using a Kruskal–Wallis test. **c,** Representative scatter plots from a bladder cancer patient in the training cohort showing associations between RNA-seq gene expression and cfChIP-seq fragment frequencies within the 280–320 bp (top) and 160–200 bp (bottom) fragment length ranges. Dashed lines depict linear regression weighted by a 2D kernel density estimate (R package MASS, v7.3-65). *R²* values summarize the strength of association.

**Extended Data Figure 4.**
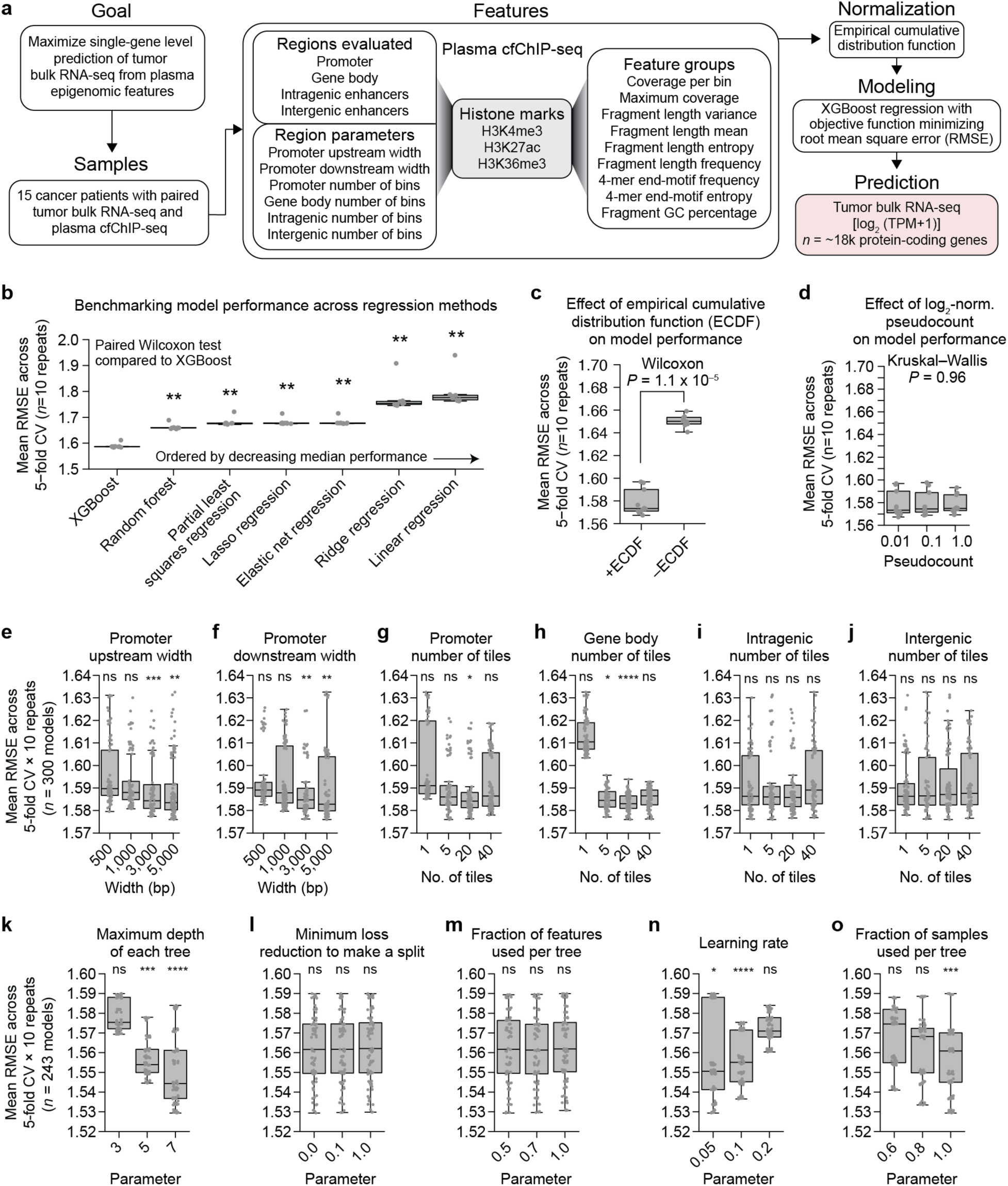
Optimization of APEX model parameters and input features. **a,** Overview of the APEX modeling framework. Paired plasma cfChIP-seq and matched tumor bulk RNA-seq from 15 cancer patients were used to train a generalized model to predict log₂(TPM+1) expression for ∼18,000 protein-coding genes. Input features included coverage metrics (per-bin and maximum), fragment-length statistics (mean, variance, entropy, and binned frequencies), 4-mer end-motif frequency and entropy, and fragment GC content. Six region-specific parameters, including window size and number of tiles, were varied as shown in panels **e–j.** Models were constructed using extreme gradient boosting regression (XGBoost; panel **b**) following empirical cumulative distribution function (ECDF) normalization (panel **c**). Log₂ pseudocount settings are shown in panel **d**. XGBoost hyperparameters were varied as shown in panels **k–o**. **b,** Benchmarking regression models. Mean root-mean-square error (RMSE) across five-fold cross-validation × 10 repeats (∼270k gene–sample pairs) for candidate models. Statistical comparisons were performed using paired two-sided Wilcoxon tests versus XGBoost. **c,** Effect of ECDF normalization. RMSE across five-fold cross-validation × 10 repeats with and without ECDF normalization. Statistical comparison performed using a paired two-sided Wilcoxon test. **d,** Effect of log₂ pseudocount choice. RMSE across five-fold cross-validation × 10 repeats using differing pseudocount values in the log₂ transformation. Statistical comparison was performed using a Kruskal–Wallis test. **e–j,** Optimization of genomic tiling and promoter windowing parameters. RMSE across five-fold cross-validation × 10 repeats for models trained with different (**e**) promoter upstream window widths, (**f**) promoter downstream window widths, (**g**) numbers of promoter tiles, (**h**) numbers of gene-body tiles, (**i**) numbers of intragenic enhancer tiles, and (**j**) numbers of intergenic enhancer tiles. Statistical comparisons were performed using two-sided Wilcoxon rank-sum tests comparing each parameter setting versus all others (ns, not significant; \**P* < 0.05; \*\**P* < 0.01; \*\*\**P* < 0.001; \*\*\*\**P* < 0.0001). **k–o,** Optimization of XGBoost hyperparameters. RMSE across five-fold cross-validation × 10 repeats for models trained with different (**k**) maximum tree depths, (**l**) minimum loss reduction (gamma) thresholds, (**m**) fractions of features per tree, (**n**) learning rates (eta), and (**o**) fractions of samples per tree (subsample). Statistical comparisons were performed using two-sided Wilcoxon rank-sum tests comparing each parameter setting versus all others (ns, not significant; \**P* < 0.05; \*\*\**P* < 0.001; \*\*\*\**P* < 0.0001).

**Extended Data Figure 5.**
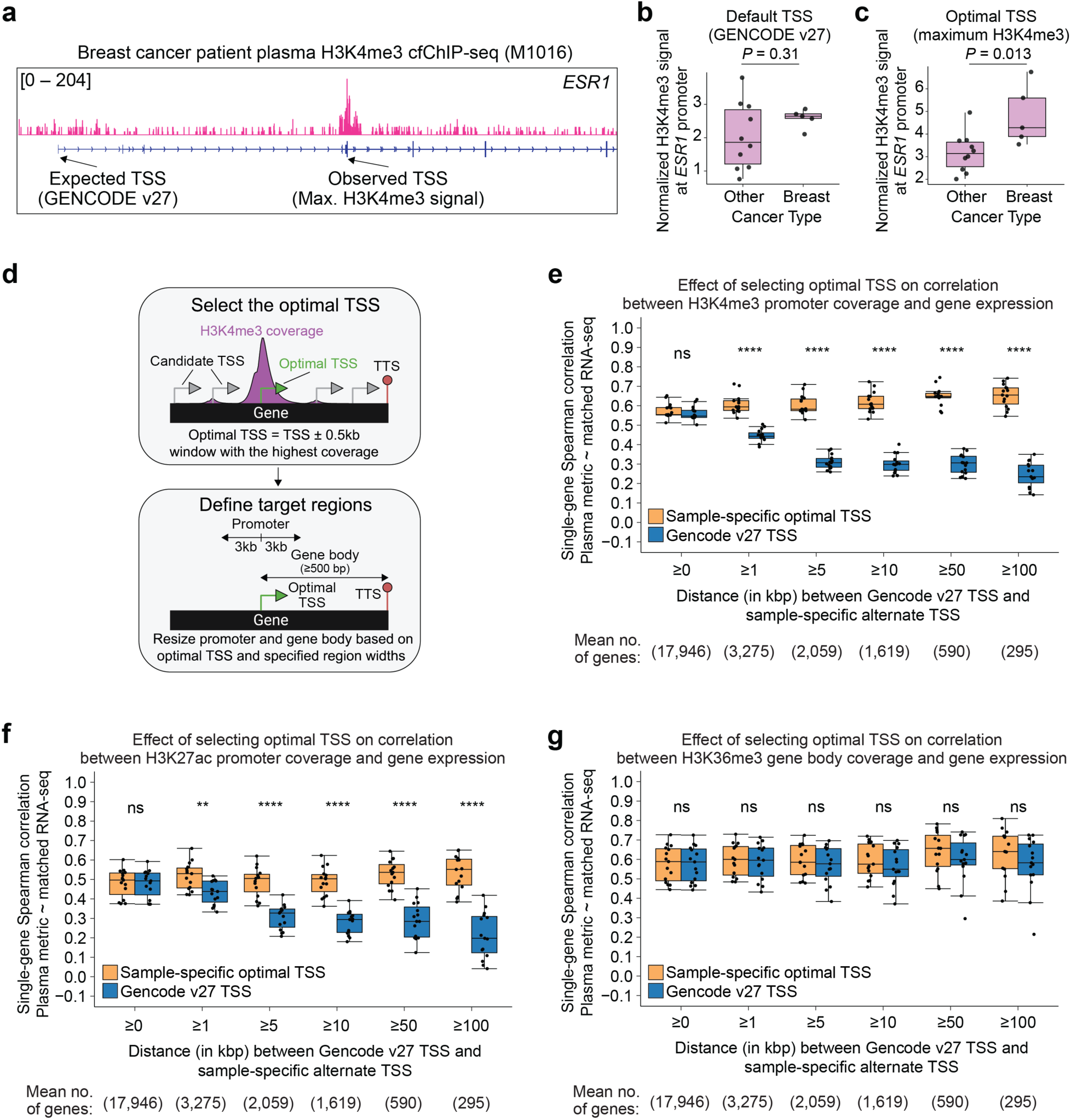
Selecting optimal transcription start sites (TSSs) improves cfChIP-seq correlation with cancer gene expression. **a,** Signal tracks of cfChIP-seq coverage for H3K4me3 in plasma from a breast cancer patient, showing the expected TSS (GENCODE v27 annotation) and the sample-specific TSS corresponding to the maximum H3K4me3 signal at the *ESR1* locus. **b,c,** Boxplots showing normalized H3K4me3 signal at the *ESR1* promoter using the default GENCODE v27 TSS (**b**) and the sample-specific optimal TSS (**c**), comparing patients with breast cancer and other cancer types in the training cohort. Statistical significance was determined using a two-sided Wilcoxon test. **d,** Schematic illustrating the method for selecting the optimal TSS and defining target regions. For each gene, the optimal TSS was identified as the ±0.5 kb window with the highest H3K4me3 coverage among all annotated TSSs. Promoter (±3 kb from TSS) and gene body (at least 500 bp) regions were then resized based on the optimal TSS and the GENCODE v27 transcriptional terminal sites (TTS). **e–g,** Boxplots showing the effect of using sample-specific optimal TSSs on correlations between plasma histone coverage and matched tumor RNA-seq in the training cohort (*n* = 15 patients) for **(e)** H3K4me3 promoter coverage**, (f)** H3K27ac promoter coverage, and **(g)** H3K36me3 gene-body coverage. Genes are stratified by the genomic distance between the GENCODE v27 TSS and the sample-specific optimal TSS. The mean number of evaluable genes across patients per group is shown below. Statistical significance was assessed using two-sided unpaired Wilcoxon tests (ns, not significant; \*\**P*< 0.01; \*\*\*\**P* < 0.0001).

**Extended Data Fig. 6.**
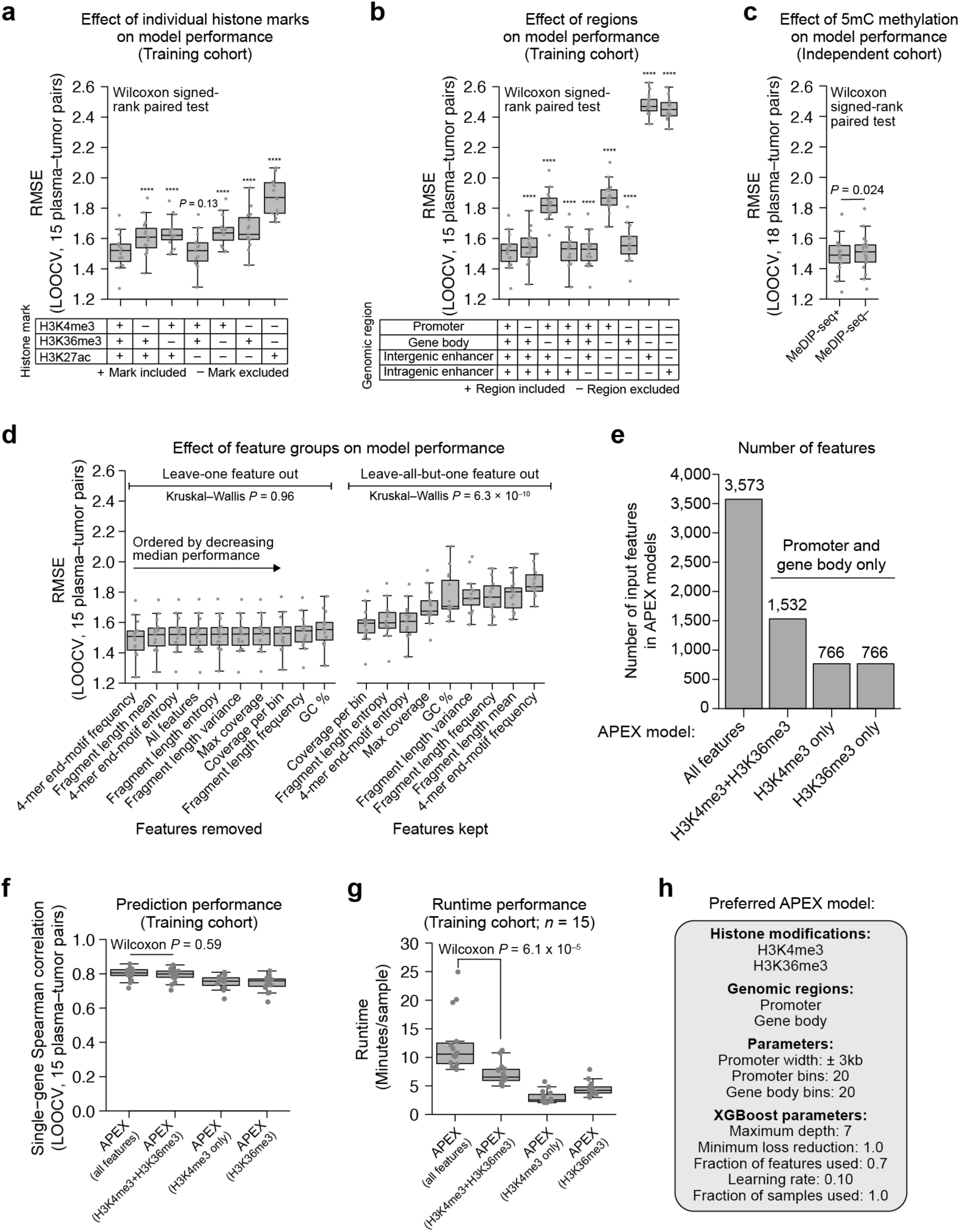
Determinants of APEX model performance and preferred feature configuration. **a,** Effect of individual histone marks on model performance. Root mean square error (RMSE) under leave-one-out cross-validation (LOOCV) across 15 plasma–tumor pairs when H3K4me3, H3K27ac, or H3K36me3 features were included or excluded. Statistical comparisons were performed using paired two-sided Wilcoxon signed-rank tests relative to the model including all marks (\*\*\*\**P* < 0.0001). **b,** Effect of genomic regions on model performance. RMSE under LOOCV across 15 plasma–tumor pairs, comparing models in which promoter, gene body, intergenic enhancer, or intragenic enhancer features were included or excluded. Statistical comparisons were performed using paired two-sided Wilcoxon signed-rank tests relative to the model including all regions (ns, not significant; \*\*\*\**P* < 0.0001). **c,** Effect of 5mC DNA methylation on model performance. RMSE under LOOCV across 18 plasma–tumor pairs using matched cfMeDIP-seq for methylation profiling (see **Methods**). Statistical comparison was performed using a paired two-sided Wilcoxon signed-rank test. **d,** Effect of feature groups on model performance. RMSE under LOOCV across 15 plasma–tumor pairs in leave-one-feature-group-out and leave-all-but-one-feature-group-in analyses. Feature groups were compared using Kruskal–Wallis tests. **e,** Number of input features across APEX model variants. Counts are shown for the full model, a refined model using H3K4me3 and H3K36me3 features from promoter and gene-body regions, and single-mark models using only H3K4me3 or H3K36me3 features. **f,** Prediction performance across APEX model variants. Single-gene Spearman correlations between predicted and measured tumor RNA-seq expression across 15 plasma–tumor pairs. Statistical comparison between full and refined models was performed using paired two-sided Wilcoxon signed-rank tests. **g,** Runtime per sample across APEX model variants. Runtime (minutes) measured on a system with 16 CPU cores and 64 GB RAM. Statistical comparison between full and refined models was performed using paired two-sided Wilcoxon signed-rank tests. **h,** Preferred APEX model configuration. Summary of histone marks, genomic regions, and XGBoost parameters selected for the final model.

**Extended Data Fig. 7.**
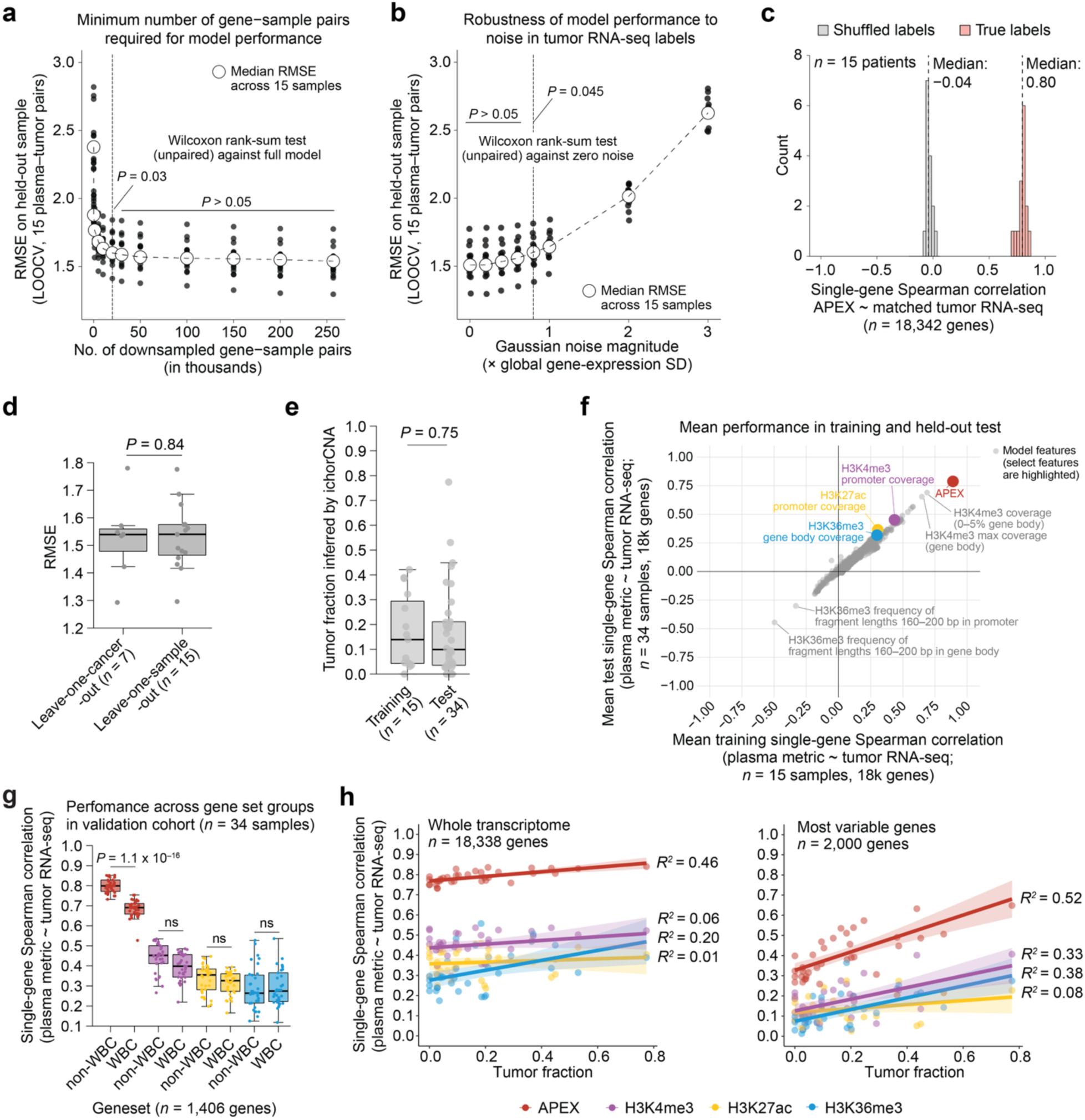
Robustness and generalization of the APEX model. **a,** Sensitivity of APEX model performance to training data size. Root mean square error (RMSE) under leave-one-out cross-validation (LOOCV) across 15 plasma–tumor pairs is shown for models trained on progressively downsampled numbers of gene–sample pairs (*x*-axis). Dashed line indicates the minimum number of feature–gene pairs required to maintain comparable performance to the full model. Statistical significance was assessed using unpaired two-sided Wilcoxon rank-sum tests against the full model. **b,** Robustness of model performance to noise in tumor RNA-seq labels. Gaussian noise (in units of the global standard deviation) was added to expression values prior to model training and RMSE was evaluated on unperturbed held-out samples using LOOCV. Statistical significance was assessed using unpaired two-sided Wilcoxon rank-sum tests against the zero-noise condition. **c,** Distribution of single-gene Spearman correlations between APEX-inferred gene expression and matched tumor RNA-seq across 18,342 genes in the training cohort (*n* = 15 patients), estimated using LOOCV. The null distribution was generated by shuffling RNA-seq labels. Median correlation values are indicated. **d,** Generalization of the APEX model to unseen cancer types via cross-validation. Comparison of RMSE between leave-one-sample-out cross-validation (*n* = 15; training on 14 samples and testing on one held-out sample) and leave-one-cancer-type-out cross-validation (*n* = 7; training on six cancer types and testing on a held-out cancer type). Statistical significance was assessed using a two-sided paired Wilcoxon rank-sum test. **e,** Tumor fraction estimated by ichorCNA in training (*n* = 15) and test (*n* = 34) cohorts. Boxplots show medians, quartiles, and 1.5 × IQR; individual samples are plotted as points. Statistical significance was assessed using a two-sided unpaired Wilcoxon rank-sum test. **f,** Mean single-gene Spearman correlation between plasma epigenomic metrics and matched tumor RNA-seq across the training (*x*-axis) and held-out test (*y*-axis) sets (*n* = 34 samples; ∼18k genes). Points represent individual feature types and colored points indicate select plasma-based metrics, including APEX-inferred gene expression and histone-mark coverage. **g,** Same as Fig. 3f, but showing single-gene Spearman correlations across WBC and non-WBC genes (1,406 genes in each set). Statistical significance was calculated by paired Wilcoxon test between WBC and non-WBC gene sets for each approach (color legend below **h**). **h,** Scatter plots show single-gene Spearman correlations between plasma-derived metrics and matched tumor RNA-seq as a function of tumor fraction. (Left) Correlations computed across the whole transcriptome (*n* = 18,338 genes). (Right) Correlations computed using the 2,000 most variable genes. Lines indicate linear regression fits with 95% confidence intervals. Coefficients of determination (*R²*) are shown for each metric.

**Extended Data Figure 8.**
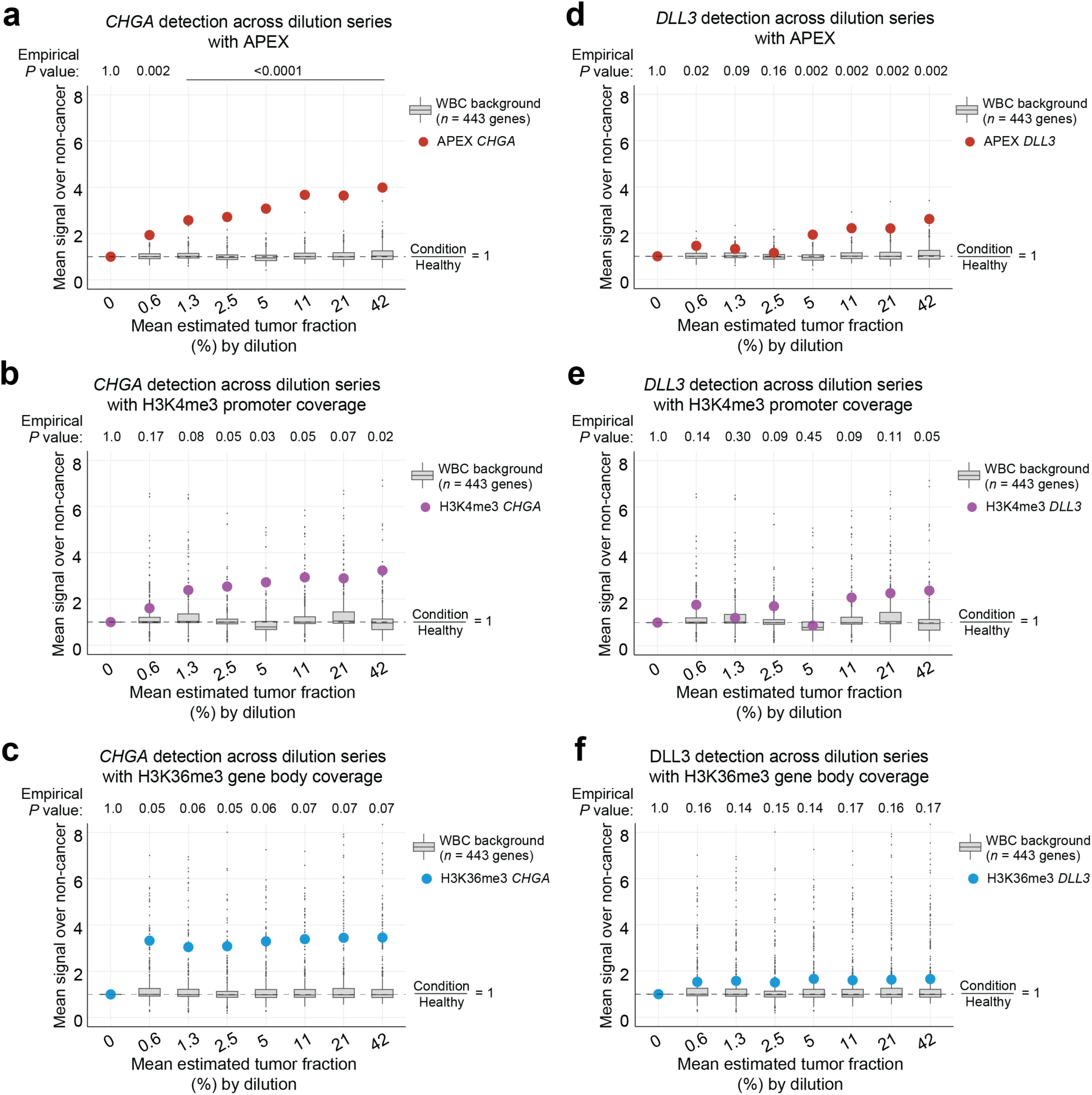
Single-gene detectability with APEX and individual cfChIP-seq across experimentally diluted tumor fractions. **a–f**, Detection of *CHGA* (lineage marker; **a–c**) and *DLL3* (therapeutic target; **d–f**) across the cfChromatin dilution series (related to Fig. 3g**–i**) using APEX, H3K4me3 promoter coverage, and H3K36me3 gene-body coverage. For each dilution, the *x*-axis reflects the mean effective tumor fraction derived from the three starting tumor fractions. The *y*-axis reflects mean signal enrichment relative to non-cancer baseline, indicated by a colored circle for the target gene, and compared with a background distribution of enrichment across 443 WBC-signature genes (gray points and boxplots) to derive empirical *P-*values. Boxplots show medians, quartiles, and 1.5 × IQR.

**Extended Data Figure 9.**
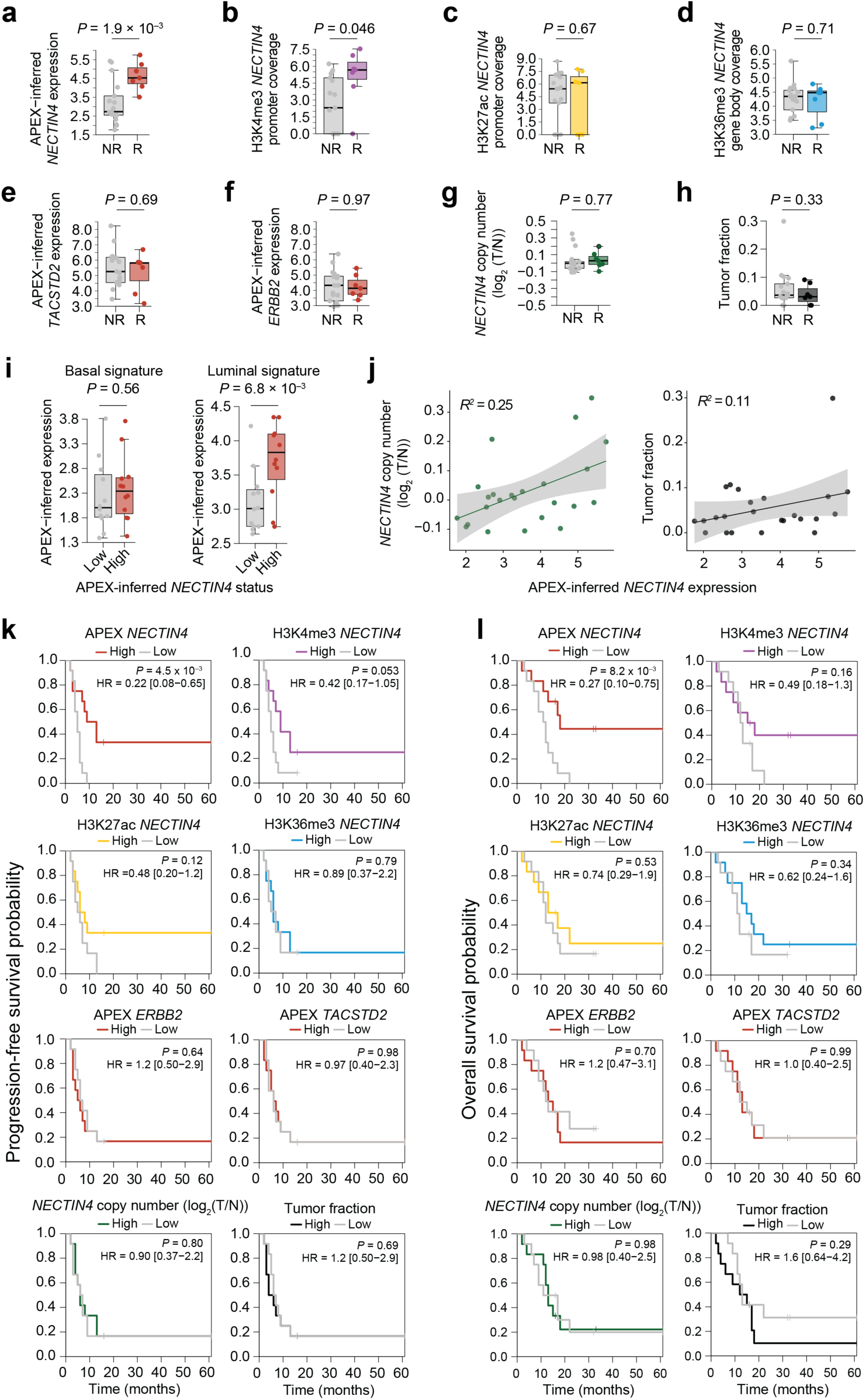
Baseline plasma-based epigenomic and genomic features and clinical outcomes in metastatic bladder cancer treated with enfortumab vedotin. **a–d,** Comparison of baseline plasma-inferred *NECTIN4* expression between non-responders (NR; *n* = 17) and responders (R; *n* = 7) to enfortumab vedotin (EV). Responders were defined as patients with complete or partial response (CR/PR), and non-responders as those with stable or progressive disease (SD/PD). *NECTIN4* expression was quantified using **(a)** APEX-inferred expression, **(b)** H3K4me3 promoter coverage, **(c)** H3K27ac promoter coverage, and **(d)** H3K36me3 gene-body coverage. **e–h**, Comparison of additional antibody-drug conjugate (ADC) targets and genomic features between NR and R, including **(e)** APEX-inferred *TACSTD2* (TROP2) expression, **(f)** APEX-inferred *ERBB2* expression, **(g)** plasma-inferred *NECTIN4* copy number variation (chr1:161–161.5 Mb, hg19; log₂ tumor/normal), and **(h)** plasma tumor fraction estimated using ichorCNA (v0.2.0). **i,** Association of APEX-inferred *NECTIN4* expression, dichotomized into high and low groups by the cohort median, with basal and luminal urothelial lineage signatures. **j,** Correlation of APEX-inferred *NECTIN4* expression with plasma-inferred *NECTIN4* copy number variation (chr1:161–161.5 Mb, hg19; log₂ tumor/normal) (left) and plasma tumor fraction estimated using ichorCNA (v0.2.0) (right). Linear regression fits are shown with coefficients of determination (*R²*). Shaded regions denote 95% confidence intervals. **k,** Progression-free survival and **l,** overall survival stratified by high versus low levels of each indicated baseline plasma-derived feature, with high and low groups defined by median dichotomization of the corresponding feature across the cohort. Features shown include histone mark-specific cfChIP-seq coverage at the *NECTIN4* locus (H3K4me3 promoter, H3K27ac promoter, H3K36me3 gene body), APEX-inferred *NECTIN4* expression, APEX-inferred expression of additional ADC targets (*TACSTD2*, *ERBB2*), plasma-inferred *NECTIN4* copy number, and plasma tumor fraction. Survival curves were compared using two-sided log-rank tests; hazard ratios (HR) and 95% confidence intervals are shown. Box plots indicate the median and interquartile range; whiskers denote 1.5 × IQR. For **a–i**, *P*-values were calculated using unpaired two-sided *t*-tests.

**Extended Data Figure 10.**
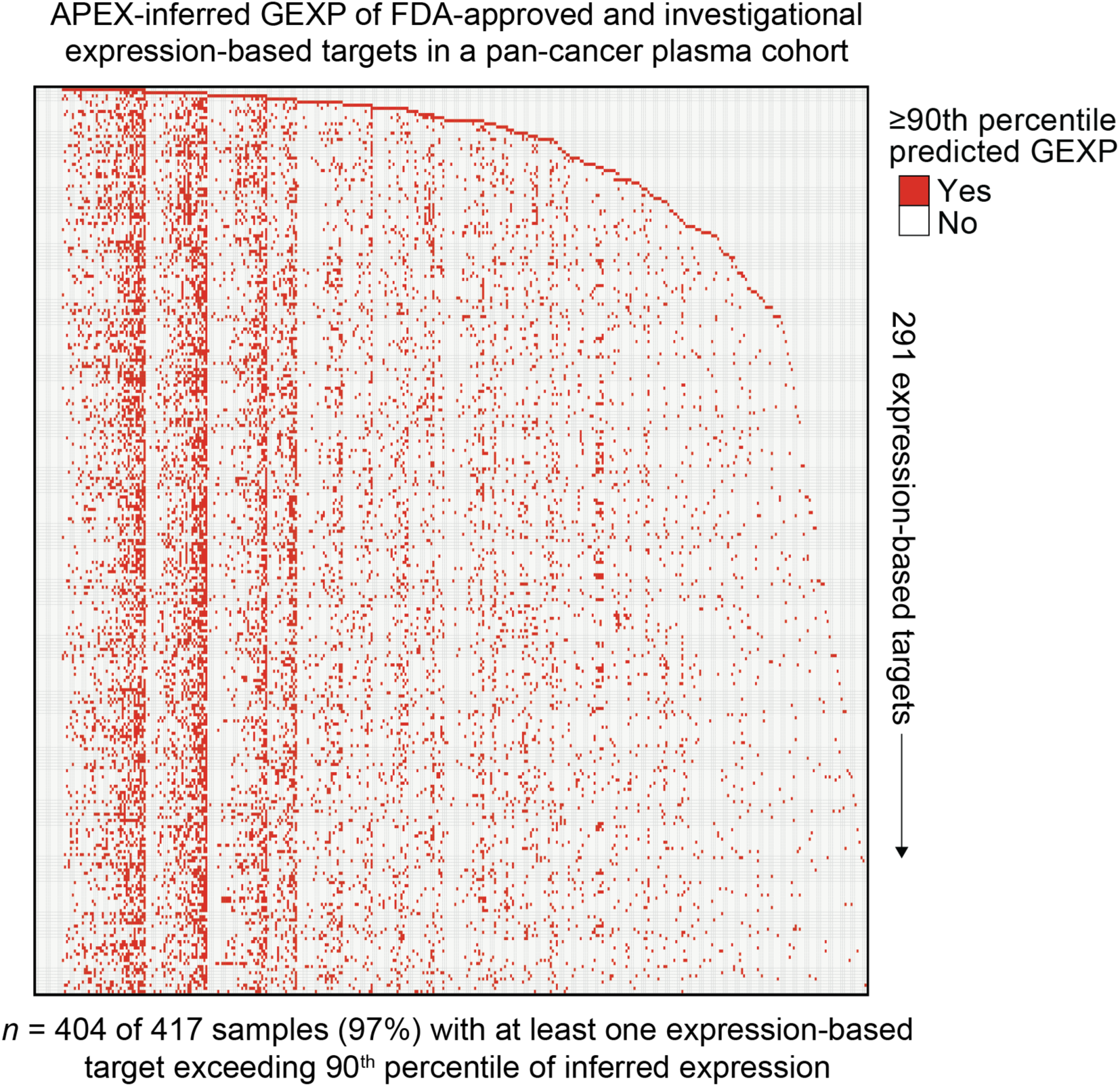
APEX-inferred expression of cancer targets in a pan-solid-cancer plasma cohort. Heatmap showing APEX-inferred gene expression (GEXP) of FDA-approved or investigational cancer targets (*n* = 291) for solid tumors across 417 plasma samples. Red indicates samples with predicted expression at or above the 90th percentile for a given target. Samples (columns) are ordered first by the dominant inferred cancer target in each sample and then secondarily by the number of cancer targets exceeding the 90th percentile of inferred expression.

## METHODS

### 1. Ethical compliance

All processing of plasma samples for cfDNA and cfChIP-seq assays was performed under Dana-Farber/Harvard Cancer Center IRB protocol 21-764, in accordance with institutional guidelines, HIPAA regulations, and the Declaration of Helsinki. For all clinical studies, patients provided written informed consent for genomic profiling and data sharing for research purposes. Plasma and tissue samples were collected under IRB-approved protocols for metastatic pancreatic cancer (DFCI 03-189) and metastatic urothelial cancer (DFCI 02-021) patients. IRB approvals for the previously published Nature Medicine study and the MOSCATO trial are described in the original publications. Additional cohort-specific details are provided in the respective Methods sections and **Supplementary Tables 3, 7, 10, 11, and 12.**

### 2. MOSCATO clinical cohort for model training and independent validation

#### 2.1. Clinichal design and patient selection

Patients aged 18 years or older with unresectable or metastatic cancer, who had progressed after at least one prior line of therapy or had limited treatment options and at least one accessible biopsy site, were enrolled in the Molecular Screening for Cancer Treatment Optimization (MOSCATO) clinical trial (NCT01566019), a monocentric, prospective study conducted at Gustave Roussy^20^. The study consisted of two sequential phases: MOSCATO–01 (2011–2016) and MOSCATO–02 (2017–2020). Patients from MOSCATO–01 with matched plasma and tumor RNA-seq data were assigned to the training cohort, whereas those from MOSCATO–02 were assigned to the test cohort. All participants provided written informed consent for trial participation, genomic and transcriptomic profiling, and data sharing for research purposes. The protocol was approved by the Institutional Review Board of Gustave Roussy and conducted in accordance with the Declaration of Helsinki and Good Clinical Practice guidelines.

#### 2.2. Plasma sample collection and processing

Plasma samples from the MOSCATO–01 and MOSCATO–02 clinical trials were collected on the same day as the tissue biopsy, corresponding to the participant’s inclusion date in the clinical trial^20^. Blood samples (3–10 mL) were drawn into EDTA tubes (BD Vacutainer, Beckton Dickinson and Company), then centrifuged at 1,000 × *g* for 10 minutes within 4 hours of collection. The resulting supernatant, containing the plasma, was subsequently centrifuged at 14,000 *× g* for 10 minutes at room temperature. The plasma was then stored at −80°C until further analysis.

#### 2.3. Tumor biopsy collection and pathological assessment

Needle biopsies were obtained as described in Pradat et al., *Cancer Discovery,* 2023^20^ from a metastatic lesion and immediately frozen in liquid nitrogen. A senior pathologist evaluated tumor cellularity on the hematoxylin and eosin-stained slide from the biopsy used for DNA and RNA extraction, which are reported in **Supplementary Tables 1 and 7**.

#### 2.4. RNA library preparation and sequencing

Tumor RNA was extracted as described in Pradat et al., *Cancer Discovery,* 2023^20^, using the AllPrep DNA/RNA Mini Kit (QIAGEN, 80204) following the manufacturer’s instructions. RNA libraries were then prepared using the TruSeq Stranded mRNA kit, adhering to the supplier’s guidelines. PolyA-containing mRNA molecules were purified from 1 μg of total RNA using oligo(dT)-coupled magnetic beads with the Magnetic mRNA Isolation Kit (NEB, S1550S). RNA fragmentation was performed using divalent cations at elevated temperatures to generate 300 to 400 bp fragments. This was followed by double-strand cDNA synthesis, Illumina adapter ligation, and PCR amplification of the cDNA library for sequencing. Paired-end sequencing was conducted on the Illumina NextSeq 500 (75 bp).

#### 2.5. RNA-sequencing processing and gene expression quantification

Tumor RNA-sequencing data were processed as described in Pradat et al., *Cancer Discovery*, 2023^20^, using the reproducible Snakemake workflow available at https://github.com/gustaveroussy/MetaPRISM_Public, which implements the pipeline described in https://github.com/gevaertlab/RNASeq_pipeline. Raw FASTQ files were preprocessed with Trim Galore (v0.4.4)^39^ to remove adapter sequences and low-quality bases. Transcript-level quantification was performed with Kallisto (v0.44.0)^40^ using the GENCODE v27 reference transcriptome (FASTA and GTF files; ftp.ebi.ac.uk/pub/databases/gencode/Gencode_human/release_27/).

Transcript-level estimates were summarized to gene-level counts using TxImport (v1.16.0)^41^, yielding expression estimates for 58,288 Ensembl genes. Gene identifiers were converted to gene symbols using the GENCODE v27 GTF annotation (GRCh37 lift-over), and multiple transcript entries mapping to the same gene were aggregated by summation. After filtering to genes with non-zero expression across all samples, 18,351 genes remained for analysis.

To account for sequencing depth and gene length, raw read counts were normalized to transcripts per kilobase million (TPM).

The TPM for gene *j* in sample *k* was calculated as:

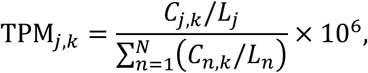

where *C_j,k_* is the raw read count for gene *j* in sample *k*, *L_j_* is the gene length in kilobases (kb), and *N* is the total number of genes quantified in that sample. To stabilize variance for downstream comparative analyses, these values were log₂-transformed using a pseudocount of 1:

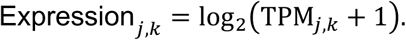

### 3. Cell-free chromatin immunoprecipitation sequencing (cfChIP-seq)

Antibodies were covalently conjugated to Dynabeads M-270 Epoxy (Invitrogen, 14302D) using the Dynabeads Antibody Coupling Kit (Invitrogen, 14311D) according to the manufacturer’s instructions, with 5 µg of antibody coupled to 1 mg of beads for at least 16 h at 37 °C with rotation. After coupling, beads were stored at 50 µg ml⁻¹ at 4 °C and pre-cleared in 0.1% BSA in PBS for 5 min at 4 °C prior to use. For chromatin immunoprecipitation, 810 µl of thawed plasma was supplemented with protease inhibitor cocktail (Roche, 11873580001), centrifuged at 3,000g for 15 min at 4 °C to remove residual debris, and combined with 90 µl of salt–detergent buffer to solubilize cell-free chromatin. The resulting 900 µl of plasma was incubated with 1 µg of antibody-coupled magnetic beads, with sequential immunoprecipitation against H3K27ac (6 h), H3K36me3 (overnight), and H3K4me3 (3 h) performed on the same plasma sample at 4 °C with rotation; for gene-body benchmarking analyses, all gene body–directed antibodies were incubated overnight. Antibody details are provided in **Supplementary Table 2**. Immunoprecipitated chromatin was preserved in chromatin-preserving buffer (50 mM Tris-HCl pH 7.6, 1 mM CaCl₂, 0.2% NP-40, 5 mM sodium butyrate, complete protease inhibitor cocktail, and 0.5 mM phenylmethylsulfonyl fluoride) prior to washing. Beads were washed sequentially three times each with 150 µl of low-salt wash buffer (0.1% SDS, 1% Triton X-100, 2 mM EDTA, 150 mM NaCl, 20 mM Tris-HCl pH 7.5), 150 µl of high-salt wash buffer (500 mM NaCl), and 150 µl of LiCl wash buffer (250 mM LiCl, 1% NP-40, 1% sodium deoxycholate, 1 mM EDTA, 10 mM Tris-HCl pH 7.5), followed by a final rinse in TE buffer. Chromatin was eluted and reverse-crosslinked in DNA extraction buffer (0.1 M NaHCO₃, 1% SDS) supplemented with Proteinase K (0.6 mg ml⁻¹; QIAGEN, 19131) and RNase A (0.4 mg ml⁻¹; Thermo Fisher Scientific, 12091021) by sequential incubation at 37 °C for 10 min, 50 °C for 1 h, and 65 °C for 90 min. DNA was purified using the ChIP DNA Clean & Concentrator kit (Zymo Research, #D5205). Sequencing libraries were prepared using the ThruPLEX DNA-Seq kit (Takara Bio, R400675), purified with AMPure XP beads (Beckman Coulter, A63880), assessed for size distribution using an Agilent 2100 Bioanalyzer with a High Sensitivity DNA chip (Agilent, 5067-4626), and sequenced as 150-bp paired-end reads on the Illumina NovaSeq X Plus platform (Novogene).

### 4. Assessment of DNA methylation features for plasma-based gene expression

Methylated DNA immunoprecipitation and sequencing (cfMeDIP-seq) was performed as previously described to assess whether genome-wide 5-methylcytosine profiles could further improve APEX performance^16,42^. We generated cfMeDIP-seq profiles for 18 samples from the prospective validation cohort, each of which already had matched cfChIP-seq data for H3K4me3, H3K27ac and H3K36me3. To quantify the incremental predictive value of methylation, we trained APEX models with and without cfMeDIP-derived features and evaluated performance using leave-one-out cross-validation across these 18 samples. Inclusion of cfMeDIP-seq features yielded a statistically significant but modest improvement in RMSE relative to the model without cfMeDIP-seq (unpaired Wilcoxon test, *P* = 0.024; median RMSE: 1.49 vs. 1.51). Given the small effect size, additional cost, and increased experimental complexity associated with cfMeDIP-seq, methylation features were not incorporated into the final APEX model (**Methods,** “Development of APEX”).

### 5. Cell-free DNA isolation and low-pass whole genome sequencing (LP-WGS)

Cell-free DNA was isolated from 1 mL of plasma using the Circulating Nucleic Acids Kit (QIAGEN, 55114), eluted in AE buffer, and stored at −80 °C. Sequencing libraries were prepared using the ThruPLEX DNA-Seq kit (Takara Bio, R400675) according to the manufacturer’s instructions and purified with AMPure XP beads (Beckman Coulter, A63880). Libraries were sequenced on an Illumina NovaSeq X Plus system (Novogene) with 150-bp paired-end reads at approximately 1–2× genome-wide coverage. Copy-number profiles and plasma tumor fraction were inferred using the ichorCNA R package (v0.2.0)^21^, which estimates tumor-derived DNA content based on read depth across genomic bins using default parameters.

### 6. Processing of cell-free ChIP-seq and DNA sequencing data

FASTQ files were demultiplexed with bcl2fastq2 (RRID:SCR_015058; v2.19) to generate sample-specific files based on unique barcode sequences. All subsequent preprocessing steps were performed using a reproducible Nextflow-based pipeline (SNAPIE v1.5.1) optimized for cfChIP-seq analysis (https://github.com/prc992/SNAPIE)^43^. Briefly, paired-end sequencing reads were adapter-trimmed using Trim Galore (v0.6.10)^39^ (trim_method set to TRIM_GALORE), but quality trimming was disabled (--quality 0) to preserve fragment length distributions and end-motif sequences. Trimmed reads were aligned to the hg19 reference genome (GCA_000001405.1) using Burrows-Wheeler Aligner (v0.7.18)^44^ with extended insert size modeling (–I 250,200,2000,10) to prevent misclassification of short and long cfDNA fragments as improperly paired. Mapped reads were filtered to retain uniquely aligned reads and sorted by coordinates using samtools (v1.15.1). PCR and optical duplicates were removed using Picard MarkDuplicates (v2.27.4; RRID:SCR_006525). Low-quality reads were removed using samtools (v1.15.1) with flags –f 3 –F 3844 –q 30 to retain only properly paired, high-confidence (mapping quality ≥ 30) reads, while excluding unmapped, secondary, QC-failed, duplicated, and supplementary reads.

The resulting BAM files were converted into BEDPE format using BEDtools (v2.30.0)^45^ bamtobed with the –bedpe flag and fragments with non-negative lengths were retained. Fragments overlapping with regions in the ENCODE hg19 exclusion list (ENCFF001TDO) were removed. The resulting fragment files from SNAPIE were imported in R as GRanges objects (GenomicRanges v1.56.2)^46^ and extremely long (>500 bp) and short (<20 bp) fragments were removed for downstream analysis. Coverage tracks were generated from aligned BAM files using bamCoverage (deepTools v3.5.6) with a 10-bp bin size and RPKM normalization, and exported in bigWig format.

### 7. Enrichment score calculation and quality control

To assess signal-to-noise quality of cfChIP-seq libraries, we computed enrichment scores by comparing chromatin signal over genomic regions where each histone modification is expected to occur (“on-target”) versus regions where it should be depleted (“off-target”). Genome-wide 18-state ChromHMM annotations across 833 biosamples were obtained from the Epimap project (hg19; (https://personal.broadinstitute.org/cboix/epimap/ChromHMM/observed_aux_18_hg19/CALLS/)^47^) and partitioned into 200-bp bins. For each histone mark, on-target regions were defined as bins annotated with mark-associated ChromHMM states in >50% of biosamples (H3K27ac: 1_TssA, 3_TssFlnkU, 8_EnhG2, 9_EnhA1; H3K4me3: 1_TssA, 2_TssFlnk, 3_TssFlnkU, 4_TssFlnkD, 8_EnhG2, 14_TssBiv; H3K36me3: 5_Tx, 7_EnhG1, 8_EnhG2). Off-target regions were defined as bins lacking any corresponding on-target ChromHMM states. ENCODE blacklist regions (https://github.com/Boyle-Lab/Blacklist/blob/master/lists/hg19-blacklist.v2.bed.gz)^48^, chrX, and chrY were excluded, and off-target regions within 10 kb of on-target regions were removed. Remaining regions were merged and filtered to retain intervals ≥1 kb, yielding 11,530/219,810 (H3K27ac), 13,795/281,427 (H3K4me3), and 12,721/90,269 (H3K36me3) on-/off-target regions, respectively. To quantify immunoprecipitation (IP) efficiency and background noise, we calculated enrichment scores as the ratio of normalized fragment coverage in on-target versus off-target regions. For each sample and histone mark, the enrichment score was defined as:

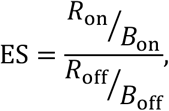

where *R*_on_ and *R*_off_ represent fragment counts in on-target and off-target regions, respectively, while *B*_on_ and *B*_off_ denote the corresponding genomic lengths.

For the training and test cohort (MOSCATO–1 and MOSCATO–2 cohorts), samples were required to meet quality thresholds of enrichment score >7 H3K4me3 and H3K27ac, and >2 for H3K36me3, with a minimum of 1 million uniquely mapped fragments per sample. For all other cohorts, no strict quality threshold was applied unless otherwise stated.

### 8. Reference epigenomic analyses and benchmarking of gene-body histone mark antibodies

#### 8.1. Calculating percent of human genome marked by histone modifications

To assess genome-wide coverage of gene body histone marks, we analyzed publicly available ChIP-seq datasets (*n* = 56) from the Roadmap Epigenomics Project^49^. These included H3K36me3 (*n* = 56), H3K79me1 (*n* = 5), H3K79me2 (*n* = 10), H3K9me1 (*n* = 3), and H4K20me1 (*n* = 11), with paired H3K4me3 (*n* = 56) and H3K27ac (*n* = 48) profiles for comparison (**Extended Data Fig. 1a**). Narrow peak calls for H3K4me3 and H3K27ac and broad peak calls for the gene body marks were downloaded from the Roadmap data portal (https://egg2.wustl.edu/roadmap/web_portal/processed_data.html). The hg19 reference genome coordinates were obtained using the BSgenome.Hsapiens.UCSC.hg19 R package (v1.4.3). To ensure mappability, regions prone to artifacts, including the ENCODE hg19 blacklist (ENCFF001TDO), genomic gaps (e.g., centromeres, telomeres), fix patches, and alternative haplotypes from the ‘Feb. 2009 GRCh37/hg19’ assembly, were excluded. Genome-wide coverage for each histone mark was calculated as the total width of non-overlapping peaks divided by the mappable genome size (2.92 × 10⁹ bp).

#### 8.2. Association between histone ChIP-seq and RNA-seq

Publicly available ChIP-seq and paired RNA-seq datasets (*n* = 56) from the Roadmap Epigenomics Project^49^ were downloaded from https://egg2.wustl.edu/roadmap/web_portal/processed_data.html and analyzed to assess the relationship between histone marks at promoters, intergenic enhancers, intragenic enhancers, and gene bodies and RNA transcription (**Extended Data Fig. 1b**). Promoter regions were defined as ±3 kb from the transcription start site (TSS) using GENCODE v27 annotation lifted to the hg19. Gene body regions were defined as the region from the TSS to the transcription termination site (TTS). Enhancer regions were identified using the GeneHancer database (v5.22)^22^, selecting only high-confidence enhancers (regulatory_element_type = “Enhancer” and is_elite = 1). Coverage was computed by counting the number of aligned sequencing reads from the ChIP-seq tagAlign files that overlapped each defined region (promoter, gene body, or enhancers) using the countOverlaps function in GenomicRanges (R v1.56.2)^46^. For enhancers (intergenic and intragenic), coverage was averaged across all associated enhancers, weighted by the element–gene association score (combined_score).

ChIP-seq coverage was normalized to account for region length, e.g. gene-body signal by gene length, enhancer signal by the mean length of associated enhancers weighted by their GeneHancer combined_score, and promoter signal by a fixed width of 6 kb centered on the TSS. RNA-seq read counts (“57epigenomes.N.pc.GENESYMBOL.txt”; sample “E000”, representing Universal Human Reference RNA, was excluded) were normalized to transcripts per million (TPM) by dividing raw counts by gene length, scaling by the total read count per sample, and multiplying by one million. Only genes annotated in all relevant datasets (e.g. RNA-seq and GeneHancer v5.22 for enhancers; RNA-seq and GENCODE v27 annotations for promoters and gene bodies) were included in the analysis. Spearman correlations were computed between histone mark signals at promoters, gene bodies, and enhancers and TPM-normalized gene expression.

#### 8.3. Benchmarking gene body histone mark antibodies for cfChIP-seq

We benchmarked antibodies targeting gene body-associated histone modifications to assess their utility for cell-free chromatin immunoprecipitation sequencing (cfChIP-seq) in plasma. Antibodies against H3K36me3, H4K20me1, H3K9me1, H3K79me1, H3K79me2, and H3K27me1 were selected based on prior usage in the literature, commercial availability, and reported validation (**Supplementary Table 2**). Antibodies were conjugated to epoxy magnetic beads and cfChIP-seq was performed as described above in “Cell-free chromatin immunoprecipitation (cfChIP-seq).” Sequenced libraries were processed as described in “Processing of cell-free ChIP-seq and DNA sequencing data”. Read-level coverage over regions of interest was calculated by counting aligned fragments overlapping gene body regions using the countOverlaps function in GenomicRanges (R v1.56.2)^46^, normalized for gene length. As a positive control, each antibody was also tested in 1 µg fixed LNCaP chromatin, which was prepared as previously described.

We benchmarked antibody performance according to three criteria:

1. Number of unique fragments: The number of deduplicated, uniquely aligned fragments recovered from cfChIP-seq libraries after sequencing.
2. Gene body enrichment score: To assess immunoprecipitation specificity, we calculated the enrichment of recovered fragments in gene-body associated chromatin states (“Tx”, “EnhG1”, “EnhG2”) as described in “Enrichment score calculation and quality control”. As a negative control, we used enrichment estimates from low-pass whole-genome sequencing (WGS) data from the 15 samples in the training cohort, which are expected to show minimal gene body coverage bias (expected enrichment score ≈ 1).
3. Prostate-cancer-specific signal recovery: To evaluate signal specificity for disease-relevant transcription, we compared cfChIP-seq gene-body signal in plasma from patients with prostate cancer to that from non-cancer donors. For each sample, we computed the mean log₂-normalized gene-body signal (normalized by gene length) across the top 500 prostate-specific genes and subtracted the mean log₂-normalized signal across the top 500 white blood cell (WBC)-specific genes (see “Defining cancer- and normal-subtype-specific gene signatures” and **Supplementary Table 8**). This difference corresponds to the log₂ ratio of prostate-selective cfChIP-seq signal. We compared these prostate-selective values between antibody- and operator-matched prostate cancer and non-cancer plasma samples to identify antibodies that maximized recovery of tumor-specific signal.

### 9. Defining cancer- and normal-subtype-specific gene signatures

#### 9.1. Data acquisition and preprocessing

RNA-seq expression data for The Cancer Genome Atlas (TCGA) and Genotype-Tissue Expression (GTEx) projects were obtained from the UCSC Xena Browser (https://xenabrowser.net/datapages/?cohort=TCGA%20TARGET%20GTEx; accessed June 7, 2025). The datasets included 10,535 TCGA tumor samples and 7,792 GTEx normal samples, provided in transcripts per million (TPM) units with matched phenotype annotations. Additional clinical and histopathologic annotations were retrieved using the R package TCGABiolinks (v2.36.0)^50^. These metadata were used to further subclassify breast cancers into ER-positive or ER-negative groups based on the er_status_by_ihc field, and esophageal cancers into adenocarcinoma or squamous cell carcinoma subtypes based on the primary_diagnosis field.

RNA-seq data for small-cell lung cancer (SCLC; *n* = 81) were obtained in log(FPKM+ 1) format from Supplementary Table 10 of George *et al.*, *Nature*, 2015^51^. Values were unlogged, the pseudocount of 1 was subtracted, and the remaining FPKM values were converted to TPM by dividing each gene’s FPKM by the total FPKM per sample and multiplying by one million. To supplement the largely localized prostate cancers represented in TCGA, RNA-seq data (in TPM units) for metastatic prostate cancer subtypes, neuroendocrine prostate cancer (NEPC; *n* = 15) and androgen receptor–pathway active prostate cancer (ARPC; *n* = 129), were obtained from the “Prostate Cancer Atlas” (https://www.prostatecanceratlas.org/api/files/public/gc_txi.abundance.fst), specifically from the “WCDT-MCRPC”^52^ and “T.C.B. 2015” (dbGaP study accession: phs000909.v1.p1) datasets.

Gene symbols were harmonized across all datasets using the HGNChelper R package (v0.8.15)^53^. All expression matrices in TPM units were merged on intersecting gene symbols, rescaled to a total of one million transcripts per sample, and subsequently log₂(TPM + 1) transformed. The resulting unified matrix comprised 18,551 samples spanning 91 phenotypes, including 38 cancer types and 53 normal tissues.

#### 9.2. Derivation of cancer- and normal-specific gene signatures

Cancer- and normal-specific gene signatures were derived from the unified TCGA–GTEx–Prostate Cancer Atlas–SCLC dataset described above. For each cancer type, cancer-specific genes were defined as those significantly enriched (log₂ fold change > 1; *P-*value adjusted for multiple hypothesis testing by the Benjamini-Hochberg method < 0.01) relative to (i) all other TCGA cancer types and (ii) all normal phenotypes in GTEx. Likewise, normal-specific genes were defined by comparing each normal tissue type to (i) all other GTEx normal tissues and (ii) all TCGA cancers using the same thresholds. Up to 500 overlapping differentially expressed genes (DEGs) shared between the two respective comparisons were retained as the final signature for each tissue or cancer type.

For analyses presented in **Figure 3k** and **4a,b**, the most specific genes among the cancers examined were determined using a specificity scoring framework. For each gene, the mean expression within each sample group was first computed. The specificity score for a given gene *j* in group *g* was defined as the difference between its mean expression in that group (*E*-*_j,g_*) and the maximum mean expression in all other groups:

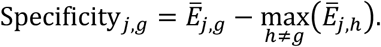

Genes were ranked within each group by their specificity scores, and the top 10 most specific genes per group were selected. The final gene list comprised the union of all group-specific top 10 genes.

All gene signatures used in this study are enumerated in **Supplementary Table 8**.

### 10. In silico screen for cell-free chromatin features associated with gene expression

To systematically identify cfChIP-seq–derived chromatin features predictive of transcriptional activity, we performed a genome-wide *in silico* correlation screen between cell-free chromatin features and matched tumor RNA-seq data (**Supplementary Table 4**).

Features were extracted as described under “Developmental of APEX: Feature extraction” (**Methods**) using default parameters. Promoter, gene body, intragenic enhancer, and intergenic enhancer regions were defined as described under “Development of APEX: Promoter, gene body, and enhancer reference” and each were divided into five equally sized bins, yielding 3,516 total features per gene. In specific instances requiring higher resolution for visualization (**Figs. 2b,c and Extended Data Fig., 2a-e)**, each genomic region was subdivided into 40 equally sized bins. Alternate promoter positions were adjusted as described under “Development of APEX: Selection of alternate promoter”.

Gene expression values were derived from log_2_(TPM + 1)-transformed RNA-seq data in the training cohort (*n* = 15). Genes with zero expression across all samples or zero evaluable fragments for any histone mark in any sample were excluded, resulting in 17,950 genes. Pairwise single-gene level Spearman correlations between each chromatin feature and gene expression value were computed using the cor() function in base R (v4.4.2) with parameters method = “spearman” and use = “pairwise.complete.obs”, such that correlations were calculated using all complete observation pairs.

### 11. Associations between chromatin fragment length and gene expression

#### 11.1. Experimental context and data sources

Associations between chromatin fragment length and gene expression were evaluated across three contexts: (1) cell-free plasma cell-free chromatin immunoprecipitated fragments, (2) non-immunoprecipitated plasma cfDNA, and (3) MNase- or Covaris-digested LNCaP chromatin. For (1), cfChIP-seq was performed using antibodies to H3K4me3, H3K27ac, and H3K36me3, with low-pass whole-genome sequencing (LPWGS) as a negative control. For (2), cfDNA from prostate (M2594; 17.5 ng) and colorectal (M2673; 20 ng) cancer patients from the MOSCATO–2 clinical trial were sequenced on the Ultima Genomics UG100 platform using XGen adapters. The mean genome coverage was 120× for M2594 and 138× for M2673 and the mean read length was 171 bp and 178 bp, respectively. For (3), ChIP-seq libraries were prepared as described under “Cell-free chromatin immunoprecipitation sequencing (cfChIP-seq)” using 0.8–1 μg of MNase-digested (“MNase-digested chromatin preparation”) or Covaris-sheared (“Covaris-sheared chromatin preparation”) LNCaP chromatin.

#### 11.2. Fragment length binning and correlation with gene expression

For each gene, fragment counts were computed across genomic regions (for example, gene bodies; **Fig. 2e**) in 10-bp bins spanning fragment lengths of 20–500 bp. Bin frequencies were normalized by the total number of fragments per region. For each fragment-length bin, Spearman correlations were computed between fragment-length frequency and matched tumor RNA-seq expression across all genes using the cor() function in base R (v4.4.2) with parameters method = “spearman” and use = “pairwise.complete.obs”, such that correlations were calculated using all complete observation pairs.

Matched bulk RNA-seq data for the cfChIP-seq dataset were obtained from the MOSCATO–1 clinical trial, whereas RNA-seq data for the deeply sequenced samples, M2594 and M2673, were obtained from the MOSCATO–2 clinical trial; all RNA-seq data were processed as described under “RNA-seq processing and quantification.” LNCaP transcript-per-million (TPM) values were obtained from the Gene Expression Omnibus (GEO; accession GSE128749) using control samples only (*n* = 3)^54^. Protein-coding transcripts were aggregated at the gene level, intersected with genes included in the promoter and gene-body reference (**Methods**, “Development of APEX: Promoter, gene body, and enhancer reference”) and averaged across control samples.

#### 11.3. MNase-digested chromatin preparation

LNCaP cells were purchased from ATCC (CRL-1740), authenticated by short tandem repeat (STR) profiling, and confirmed to match the parental LNCaP line in the ATCC database. Prior to experiments, all cells were tested for mycoplasma contamination using the LookOut Mycoplasma PCR Detection Kit (Sigma-Aldrich, #D9307). Mononucleosomal chromatin was prepared from LNCaP cells following a protocol adapted from Verzi *et al.*, *Dev. Cell*, 2010^55^. Briefly, 3 × 10⁷ cells were washed in ice-cold PBS containing protease inhibitors (Sigma-Aldrich, Cat. #11873580001), scraped, and pelleted by centrifugation (2500 × g, 3 min, room temperature). The pellet was resuspended in 900 μL digestion buffer (50 mM Tris-HCl, pH 7.6, 1 mM CaCl₂, 0.2% NP-40, 5 mM sodium butyrate, complete protease inhibitor cocktail, and 0.5 mM phenylmethylsulfonyl fluoride) and Dounce-homogenized for 1 min. After a 5-min incubation on ice, samples were warmed to 37 °C for 2 min and digested with 3 μL micrococcal nuclease (MNase; 0.2 U/μL, Sigma-Aldrich #N3755) for 7 min at 37 °C. The reaction was stopped by addition of 4.5 μL 0.5 M EDTA (final 2.5 mM). Digested chromatin was briefly sonicated for 30s and then resuspended in dilution buffer (1% NP-40, 0.5% sodium deoxycholate, 0.1% SDS, and complete protease inhibitor cocktail in PBS). Mononucleosomal enrichment (∼146 bp) was assessed using the Agilent 4150 TapeStation system and analysis software v3.2 (Agilent Technologies, Inc., Santa Clara, CA). Lysates were centrifuged (14,000 × g, 10 min, 4 °C), and the supernatant containing soluble chromatin was collected and stored at −80 °C until use.

#### 11.4. Covaris-sheared chromatin preparation

Fixed chromatin was fragmented by ultrasonication to generate random DNA fragments. Briefly, adherent LNCaP cells (1 × 10⁷) were crosslinked by directly adding formaldehyde to the culture medium to a final concentration of 1% and incubating for 10 min at room temperature. Crosslinking was quenched with 125 mM glycine for 5 min at room temperature. Cells were washed twice with ice-cold PBS containing complete protease inhibitors (Sigma-Aldrich, Cat. #11873580001), scraped, and pelleted by centrifugation (625 × g, 4 min, 4 °C). Pellets were resuspended in 1 mL dilution buffer (1% NP-40, 0.5% sodium deoxycholate, 0.1% SDS, and complete protease inhibitors in PBS) and incubated on ice with gentle rocking for 10 min. Chromatin was sheared using a Covaris AFA system (1 mL tubes; peak incident power = 140, duty factor = 5%, burst cycle = 200) for 25 min at 4 °C. Lysates were centrifuged (14,000 × g, 10 min, 4 °C), and the supernatant containing soluble chromatin was collected and stored at −80 °C until use.

### 12. Development of APEX

#### 12.1. Promoter, gene body, and enhancer reference

Protein-coding genes and their coordinates were obtained from GENCODE comprehensive gene annotation file (GRCh37, release 27) by filtering for entries where type = gene and gene_type = protein_coding. Multimapping gene entries and genes with lengths smaller than 500 bp were excluded. Promoters were defined as a symmetric 3kb window around the transcriptional start site (TSS), and gene body was defined as the region from the TSS to the transcriptional terminal site (TTS). We found that binning the promoter and gene body into 20 equally sized windows (e.g. a fixed 400 bp promoter bin size and a gene body bin size of *G_j_*⁄20, where *G_j_* denotes the gene width from TSS to TTS in bp), resulted in significantly better model performance in the training set compared to other bin sizes (**Extended Data Fig. 4g–j**).

Enhancer regions were obtained from the GeneHancer database (v5.22; GRCh37)^22^, restricting to high-confidence elements (regulatory_element_type = “Enhancer” and is_elite = 1). Original enhancer coordinates were retained. Enhancers overlapping annotated gene bodies were classified as intragenic (40,342 enhancers across 9,042 genes), whereas those located outside gene bodies were classified as intergenic (26,495 enhancers across 7,564 genes). These promoter, gene body, and enhancer annotations defined the genomic reference used for all downstream feature extraction and modeling in APEX (**Supplementary Table 5**).

#### 12.2. Selection of alternate promoter

A previous study demonstrated that transcription at protein-coding genes is regulated by context-specific promoter usage^25^. Consistent with this, we observed that histone modification patterns in plasma cfDNA mark alternative promoters in a cancer-type–dependent manner (**Extended Data Fig. 5; Supplementary Note**). To account for this variability, candidate transcription start sites (TSSs) were obtained using the proActiv R package (v1.1.18)^25^, with GENCODE GRCh37 GTF file (release 27) provided as input. Promoter coordinates were derived from the resulting PromoterAnnotation object, and the union of proActiv-defined alternative promoters and GENCODE v27 TSSs was used as the candidate TSS set for each gene (*n* = 66,976 TSSs across 18,493 protein-coding genes; **Supplementary Table 5**). For each sample, fragment coverage was quantified in a ±500 bp window around each candidate TSS, and the TSS with maximal coverage was selected as the optimal promoter. Promoter (TSS ±3 kb) and gene-body (TSS to TTS) coordinates were then updated accordingly. Optimizing transcription start site (TSS) selection in plasma epigenomic data significantly improved gene-level Spearman rank correlations with matched tumor RNA-seq. The most substantial gains were observed for genes where the selected TSS deviated by >10 kb from the default GENCODE (GRCh37, release 27) annotation (**Extended Data Fig. 5**).

#### 12.3. Feature Extraction

To develop APEX (<u>A</u>ssociating <u>P</u>lasma <u>E</u>pigenomics with e<u>X</u>pression), we extracted coverage-, size-, and sequence-based features across 18,342 protein-coding genes (**Fig. 1b,c**). Features were derived from cfDNA co-immunoprecipitated with circulating histones marked by H3K4me3, H3K27ac, or H3K36me3, and across promoter, gene body, intergenic enhancer, and intragenic enhancer regions as defined in “Development of APEX: Promoter, gene body, and enhancer reference” (**Fig. 1b, Supplementary Table 5**). The rationale and definitions for each feature are described below. Notably, for intergenic and intragenic enhancer regions, the reported metric for each gene was calculated as the weighted average across all of that gene’s associated enhancer peaks, as annotated in the GeneHancer database^22^.

#### 12.3.1. Coverage per bin

The total signal from active histone marks (H3K4me3, H3K27ac, and H3K36me3) and its distribution across promoters, enhancers (intergenic and intragenic), and gene bodies has been associated with gene expression^15^. Therefore, we quantified coverage for each bin *i* and gene *j* in region *p* as follows:

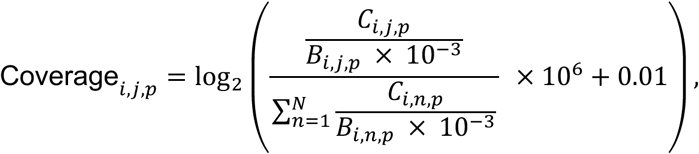

where counts (*C_i,j,p_*) represent unique fragments overlapping the bin, calculated as the number of unique fragments overlapping that bin using the findOverlaps function in the GenomicRanges R package (v1.56.2)^46^.

To account for length-related biases, counts were normalized by the bin width in kilobases, or (*B_i,j,p_*), which varied according to the number of bins, the region evaluated, and the gene considered. Specifically, promoter regions were defined as a fixed 6 kb window centered on the transcriptional start site (TSS; unless otherwise specified); gene body regions spanned the distance from the TSS to the transcription termination site (TTS), with TSS coordinates defined by Gencode v27 or adjusted to the optimal TSS as described under “Development of APEX: Selection of alternate promoter”; and intergenic or intragenic enhancer regions were defined by the weighted average (**Methods**, “Development of APEX: Promoter, gene body, and enhancer reference”) of widths corresponding to each gene’s associated enhancer peaks as annotated in the GeneHancer database^22^.

Bin-width–normalized counts were scaled by the total number of fragments (in millions) aligning to the same bin position across all genes in that sample to account for bin-specific and sample-level differences in sequencing depth. The resulting values were log₂-transformed with a small pseudocount (0.01), chosen to introduce minimal bias relative to larger pseudocounts while providing slightly improved performance in the training cohort (**Extended Data Fig. 4d**).

##### 12.3.2. Maximum coverage per region

We hypothesized that the maximal signal intensity of active histone marks, such as H3K4me3, H3K27ac, and H3K36me3, may correlate with gene expression activity. For each gene *j* and region *p*, the bin *i* with the highest coverage among the *n*_!,1_ bins was identified and included as a model feature:

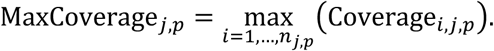

##### 12.3.3. Frequency of differently sized fragments per region

The distribution of cfDNA fragment lengths can reflect patterns of transcription factor– and nucleosome–protected DNA^56^. Fragment length frequencies have also been associated with locus-specific methylation and histone mark activity^57,58^. In our study, we identified fragment size windows that were most strongly associated with gene expression in the training cohort (**Fig. 2a,e,f**). Based on these findings, we defined five fragment size windows: 20–80 bp, 160–200 bp, 280–320 bp, 400–440 bp, and all remaining fragments shorter than 500 bp.

For each gene *j* and region *p*, the frequency of fragments within each window w was calculated as a feature in the model:

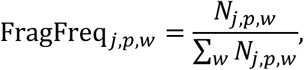

where *N_j,p,w_* is the number of fragments of size window *w* in region *p* of gene *j*, and the denominator sums the counts of fragments across all windows *w* for that same gene *j* and region *p*. This normalization produces a relative frequency for each fragment size window, allowing comparisons across genes and regions while accounting for differences in total fragment counts.

##### 12.3.4. Mean and variance of fragment length per bin

Following the rationale described in “Frequency of differently sized fragments per region”, we also computed the mean and variance of fragment lengths for each bin *i* and gene *j* in region *p*.

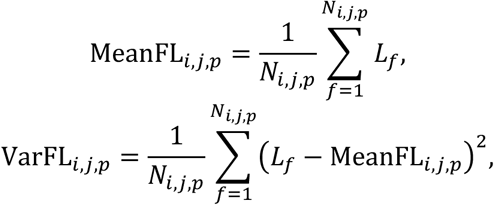

where *L_f_* is the length of fragment *f* overlapping bin *i* of gene *j* in region *p*, and *N_i,j,p_* is the total number of fragments overlapping that bin.

##### 12.3.5. Shannon entropy of fragment length per bin

Previous studies have associated fragment length diversity near the TSS with gene expression^9^. However, the relationship between the diversity of nucleosome-immunoprecipitated fragments and gene expression has not been explicitly assessed. To evaluate this, we calculated fragment length frequencies across a range of 20–500 bp, as follows:

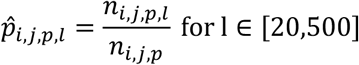

where *n_i,j,p,l_* denotes the number of fragments of length *l* in bin *i* of gene *j* and region *p*, and *n_i,j,p_* represents the total number of fragments in that bin. We then computed the Shannon entropy of these distributions within each bin *i* in gene *j* and region *p*:

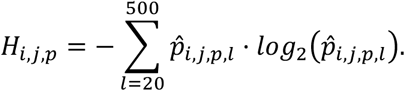

Entropy values were normalized by dividing by *log*_2_(*L*), where *L* is the total number of fragment length bins (481 for the 20–500 bp range), to normalize values between 0 and 1.

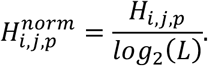

Lower entropy values indicate reduced diversity or a highly skewed fragment length distribution, while higher values reflect greater diversity or a more uniform distribution.

##### 12.3.6. Frequency of 4-mer end motifs per region

End-motif sequence biases have been associated with specific histone modifications, nucleosome positions, and tissue-specific enzymes^12,13^. To avoid potential distortions of 3′ end motifs caused by end-repair and A-tailing during double-stranded DNA library preparation, we analyzed motifs at the 5′ ends of fragments. Using bedtools getfasta (v2.30.0)^45^, we extracted 4-mer nucleotide motifs from the 5′ end of both the positive and negative fragment strands, yielding two end-motifs per fragment. For each gene *j* and 4-mer motif *m* in region *p*, the motif frequency was calculated as a relative fraction:

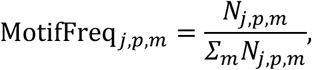

where *N_j,p,m_* is the count of motif *m* in region *p* of gene *j*, and the denominator sums the counts of all 256 possible 4-mer motifs within the same gene *j* and region *p*. This normalization produces a relative frequency for each motif, allowing comparisons across genes and regions while accounting for differences in total fragment counts.

##### 12.3.7. Shannon entropy of 4-mer end motifs per region

Several studies have suggested that the distribution of end-motif sequence biases can be associated with locus-specific epigenomic activity^12,13^. To evaluate this, we computed a motif diversity score (MDS), defined as the Shannon entropy of 4-mer frequencies for each gene *j* and region *p* across all *L* = 256 possible combinations of ‘A’, ‘C’, ‘T’, and ‘G’. For each motif *m*, the relative frequency in region *p* of gene *j* was calculated as:

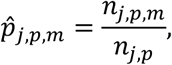

where *n_j,p,m_* is the number of fragments with motif *m* in region *p* of gene *j*, and *n_j,p_* is the total number of fragments in that region. The Shannon entropy of 4-mer end motifs (MD*S_j,p_*) for gene *j* and region *p* was then computed as:

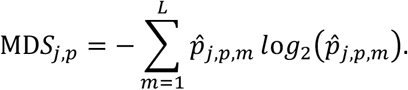

To facilitate comparisons across genes and regions, MDS values were normalized to the theoretical maximum entropy:

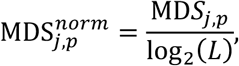

yielding values between 0 and 1. Lower MDS values indicate reduced sequence diversity or a strong preference for specific 4-mer end motifs, whereas higher values reflect greater diversity or a more uniform motif distribution.

##### 12.3.8. Average fragment GC content per bin

The GC content of DNA fragments contributes substantially influences coverage variability through both technical factors (e.g., PCR amplification bias, library preparation methods) and biological factors (e.g., promoter CpG density, transcription factor binding sites [TFBSs]). Distinguishing technical artifacts from true biological signals in ChIP-seq is particularly challenging because immunoprecipitated DNA is often enriched for GC-rich regions, especially near promoters and TFBSs. Although GC bias correction methods have been proposed for transcription factor ChIP-seq^59^, standardized approaches for histone ChIP-seq are lacking.

To capture potential GC-related effects, we incorporated the mean GC content of fragments within each bin as a feature. GC content was computed using *bedtools nuc* (v2.30.0)^45^ as the proportion of guanine and cytosine bases per fragment. For each bin *i* and gene *j* in region *p*, the mean GC content 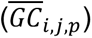 was calculated as:

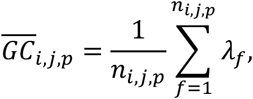

where *n_i,j,p_* is the number of fragments in that bin, and *λ_f_* is the GC fraction of fragment *f*, computed from the number of guanine (*G_f_*) and cytosine (*C_f_*) bases divided by the total fragment length (*L_f_*):

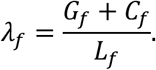

All cfChIP-seq features were compiled at the gene level to generate a unified feature matrix. For the initial analysis of the training cohort and *in silico* screen (Fig. 1–2), each annotated region was evenly tiled into five bins, yielding 3,516 parameters per sample. Features included bin-level coverage, maximum coverage per region, fragment-length statistics (Shannon entropy, mean, and variance), 4-mer end-motif entropy and frequencies, and GC content, computed for each histone modification (H3K4me3, H3K27ac, and H3K36me3). Following feature optimization (**Methods**, “Development of APEX: Feature and model optimization”), promoter and gene-body regions were divided into 20 bins, while intragenic and intergenic enhancer regions remained untiled (one bin each), resulting in 3,912 parameters.

##### 12.3.9. Feature normalization

To normalize signal intensities across samples and reduce the impact of extreme values, each feature vector was transformed to its empirical percentile distribution. An empirical cumulative distribution function (ECDF) was computed from all valid (non-missing, non-zero) values, treating zeros as missing to avoid distortion of the lower tail. For each valid observation *x*_0_, the transformed value *F*(*x*_0_) was calculated as:

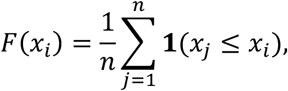

where *n* represents the number of valid observations, and **1**(·) is the indicator function, which equals 1 if the condition *x_j_* ≤ *x_i_* is met and 0 otherwise. Each valid observation was replaced by its corresponding empirical percentile. Original zero or missing values were excluded from the rank computation and retained as NA.

#### 12.4. Regression model selection

To identify the optimal statistical framework for predicting gene expression from cfChIP-seq features, we trained and compared multiple regression models in *R* (v4.4.2). In total, we evaluated seven regression frameworks: ordinary least squares (OLS), ridge, lasso, and elastic-net regression (*glmnet*, v4.1-8)^60^; partial least squares regression (*pls*, v2.8-1); random forest regression (*ranger*, v0.16.0); and gradient boosting regression (*xgboost*, v1.7.7)^61^. The input matrix comprised ECDF-normalized chromatin features (17,941 genes × 3,516 features) across 15 training samples (269,115 total gene–sample observations). Missing values for individual features were imputed with their median across all observations. The outcome variable for each gene–sample observation was log₂(TPM + 1) gene expression.

Linear regression models were fit using *glmnet* with *⍺* = 0 (ridge), *⍺* = 1 (lasso), or *⍺* = 0.5 (elastic-net). Each model was trained using 5-fold cross-validation, minimizing mean squared error (MSE), and predictive performance was summarized by the root-mean-squared error (RMSE) on held-out data. OLS regression was implemented as ridge regression with a minimal penalty (*λ* = 10^−8^). Partial least squares regression was fit with up to 30 components using segment-wise cross-validation. Tree-based models were trained with default parameters: random forests (*ranger*) with 500 trees, and gradient boosting (*xgboost*) with 500 boosting rounds and early stopping after 20 rounds without improvement.

All models were evaluated using stratified 5-fold cross-validation, repeated 10 times to ensure stability. Performance was assessed by the RMSE on held-out data. The optimal model was defined as the one yielding the lowest mean RMSE, with significance determined by a paired Wilcoxon rank-sum test (*P* < 0.05) against other models.

#### 12.5. Feature and model optimization

Feature and model parameters were jointly optimized using a two-stage grid search strategy (**Extended Data Fig. 4e–o**). In the first stage, genomic feature parameters defining the resolution of promoter, gene body, intragenic, and intergenic regions were tuned to maximize predictive performance. Candidate feature configurations were drawn from a grid specifying the number of promoter, gene body, intragenic enhancer, and intergenic enhancer tiles (1, 5, 20, or 40 each), and, for promoters, the upstream and downstream window sizes relative to the TSS (500, 1,000, 3,000, or 5,000 bp). From all possible combinations, 300 were randomly sampled for evaluation.

Each parameter set was evaluated using an XGBoost regression model in R (v.1.7.8.1; default parameters: *η* = 0.3, max_depth = 6, *γ* = 0, colsample_bytree = 1, subsample = 1, nrounds = 500, early_stopping_rounds = 20, tree_method = “hist”, objective = “reg:squarederror”, and evaluation metric = “rmse”). Model performance was assessed by five-fold cross-validation, where 15 samples were randomly partitioned into five folds. To reduce sampling bias and yield stable performance estimates, cross-validation was repeated ten times with distinct random seeds defined as *set*. *seed* = 42 + *i* for the *i*-th repeat. For each configuration, the mean RMSE across the ten repeats was recorded, and parameter importance was assessed using the Wilcoxon rank-sum test to identify the optimal feature set minimizing RMSE (**Extended Data Fig. 4e–j**).

In the second stage, XGBoost hyperparameters were optimized using the best-performing feature configuration (20 promoter tiles, 20 gene body tiles, one intragenic tile, one intergenic tile, and 3,000 bp upstream and downstream windows). A grid of 243 hyperparameter combinations was evaluated, varying learning rate (*η* ∈ {0.05, 0.1, 0.2}), maximum tree depth (max_depth ∈ {3, 5, 7}), minimum loss reduction (*γ* ∈ {0, 0.1, 1}), column subsampling rate (colsample_bytree ∈ {0.5, 0.7, 1.0}), and row subsampling rate (subsample ∈ {0.6, 0.8, 1.0}). The same 5-fold, ten-repeat cross-validation strategy was used to compute the mean RMSE for each combination, and the parameter set yielding the lowest mean RMSE was selected for final model training. Where multiple settings yielded comparable performance, parameters favoring greater regularization were selected. For example, a column subsampling rate of 0.7 was chosen despite similar RMSE at 1.0 to reduce overfitting among correlated features, and γ = 1 was selected as a conservative setting given similar performance across the tested range (**Extended Data Fig. 4k–o**).

#### 12.6. Model training

Final XGBoost regression models were trained on the complete ECDF-normalized feature matrix, comprising 3,912 features across 275,130 observations (18,342 genes × 15 samples). The target variable was log-transformed RNA-seq expression (log_2_[TPM + 1]). Training was performed using the *xgb.train* function in R with the hyperparameter set identified during model optimization.

To address varying data availability and computational constraints, we developed four versions of the APEX model: **APEX-full**, incorporating all 3,912 features; **APEX-(K4 + K36)**, a streamlined model using the most informative H3K4me3 and H3K36me3 features; and single-mark variants **APEX-K4** and **APEX-K36**. APEX-(K4 + K36) achieved performance comparable to APEX-full while being computationally faster, and APEX-K4 alone substantially improved over promoter-only (H3K4me3) coverage models. Although multi-mark models yielded the highest accuracy, single-mark APEX versions are provided for users with limited cfChIP-seq data availability.

### 13. Robustness of APEX model performance to the number of gene–sample pairs

To assess the robustness of model performance to the number of gene–sample observations, we evaluated how training sample size affects model performance. Briefly, we trained models with different gene–sample pairs in the training, using the optimized feature matrix (*n* = 1,532 features; parameters: 20 promoter and gene body tiles for H3K4me3 and H3K36me3 with promoter regions defined as ±3 kb around the TSS) and optimized XGBoost configuration (*η* = 0.1, max_depth = 7, *γ* = 1, colsample_bytree = 0.7, subsample = 1, tree_method = “hist”). Model performance was evaluated using leave-one-out cross-validation (LOOCV), where models were trained on a downsampled (e.g. 200,000, 150,000, 100,000, 50,000, 30,000, 20,000, 10,000, 5,000, 1,000, 100, and 10 observations) or full feature matrix (e.g. 256,788 gene–sample observations for 14 samples) and tested on all genes in a held-out sample (18,342 genes). Predictive accuracy was quantified as the RMSE on the held-out sample.

The minimum number of gene–sample pairs required to maintain robust performance was defined as the smallest subset size whose RMSE was not significantly higher than that of the full model, as determined by an unpaired Wilcoxon rank-sum test (*P* < 0.05). Robust performance was preserved down to ∼30,000 gene–sample pairs (**Extended Data. Fig. 7a**).

### 14. Robustness of model performance to noise in tumor RNA-seq labels

Cancer gene expression measurements can vary substantially due to differences in tumor purity and spatial heterogeneity. Moreover, multiple metastatic sites can contribute to signal in plasma cfDNA. To assess the stability of model performance under such variability, we evaluated robustness to label perturbation.

The same optimized feature matrix and hyperparameters were used, with leave-one-out cross-validation (LOOCV) as described above.

Gaussian noise was added to tumor RNA-seq labels in the training folds only (256,788 gene–sample observations across 14 samples per iteration), scaled to a specified fraction of the global standard deviation of expression values (0–10×), and truncated at zero to avoid negative values. Scaling noise to the global rather than gene-specific standard deviation ensured that perturbation magnitude was independent of intrinsic expression variability, which is correlated with biological signal strength. For each noise level (0, 0.2, 0.4, 0.6, 0.8, 1, 2, 3, 5, and 10×), models were trained on label-perturbed data and evaluated on the unperturbed held-out sample (18,342 genes). Model performance was summarized as the RMSE between predicted and observed expression.

The highest noise level at which RMSE remained statistically indistinguishable from the noise-free baseline (unpaired Wilcoxon rank-sum test, *P* < 0.05) was defined as the robustness threshold. The model remained stable up to a perturbation amplitude of approximately 0.6× the global standard deviation, indicating resilience to moderate variability in tumor RNA-seq measurements.

### 15. Evaluation of APEX generalizability across cancer types

To assess the robustness of APEX across diverse tumor contexts, we evaluated model performance using two complementary cross-cancer generalizability analyses. First, within the training cohort, we performed a leave-one-cancer-out (LOCO) procedure. The 15 training samples were grouped into seven tumor types, and for each iteration, APEX was trained on six cancer groups and evaluated on all gene–sample pairs from the excluded cancer type. This analysis quantified the extent to which models trained on a subset of cancers could predict gene expression programs in an unseen malignancy. LOCO performance was compared directly with leave-one-sample-out cross-validation (LOOCV), in which the model was trained on 14 samples and evaluated on the held-out sample. Across all iterations, no significant difference in RMSE was observed between LOCO and LOOCV, indicating that APEX performance was stable even when entire cancer types were omitted from training (**Extended Data Fig. 7d**).

To further evaluate external generalizability, we assessed APEX in an independent cohort from the MOSCATO–2 clinical trial, which comprised 34 samples spanning 18 cancer types, 12 of which were not represented in the training cohort (see “MOSCATO clinical cohort”). For each sample, Spearman correlation was computed between APEX-predicted gene expression and matched bulk tumor RNA-seq. Performance in MOSCATO–2 was comparable to that observed in the training cohort, despite differences in cohort composition, sample handling, and cancer representation (**Fig. 3f**; **Extended Data Fig. 7f**). Together, these analyses demonstrate that APEX maintains predictive accuracy across a broad range of tumor types and generalizes to malignancies not encountered during model development.

### 16. Feature contribution and importance analysis

To assess the contribution of each feature class and the relative importance of individual features to model performance, we performed systematic feature ablation and permutation-based importance analyses. For each ablation, models were retrained after excluding defined feature groups, including genomic regions (promoter, gene body, intergenic, intragenic), histone marks (H3K4me3, H3K27ac, H3K36me3), or cfDNA-derived feature categories (coverage, GC content, fragment length, and 4-mer end-motif metrics), using identical training and cross-validation procedures. Model performance was evaluated by leave-one-out cross-validation (LOOCV) and summarized as the root-mean-squared error (RMSE) between predicted and observed expression. Statistical significance of RMSE differences was determined using paired Wilcoxon signed-rank or Kruskal–Wallis tests, as appropriate.

To quantify the relative influence of individual chromatin features, we applied a model-agnostic permutation-based importance framework (DALEX v2.5.3, R)^62^. For each feature, its values were randomly permuted across samples in the final XGBoost model (trained on H3K4me3 and H3K36me3 promoter and gene body features), and the resulting degradation in model accuracy was used to estimate importance. The median loss in predictive performance across all permutations provided a stable measure of each feature’s contribution, enabling unbiased ranking of the most informative chromatin determinants of gene expression.

### 17. Benchmarking against existing fragmentomic and cfChIP-seq methods

To benchmark APEX against established fragmentomic and cfChIP-seq-based gene-expression inference methods, we selected five samples from the independent validation cohort spanning a range of tumor types, tumor fractions, and pre-library cfDNA inputs: M2187 (uterine; 19.55 ng cfDNA; ctDNA 0.45; 92× depth), M2240 (esophageal; 20 ng; ctDNA 0.37; 93×), M2246 (lung; 100 ng; ctDNA 0.77; 101×), M2331 (cholangiocarcinoma; 16.51 ng; ctDNA 0.05; 114×), and M2692 (colorectal; 32 ng; ctDNA 0.20; 117×). Sequencing depths (90–120×) were selected to match ranges commonly used in prior fragmentomic studies. For each sample, 1 mL of plasma was independently profiled by cfChIP-seq for H3K4me3, H3K27ac, and H3K36me3 using the workflow described above.

APEX predictions were generated as described above. H3K4me3 and H3K27ac promoter signals were quantified within ±3 kb of GENCODE v27 transcription start sites and normalized to DNase I hypersensitive site (DHS) coverage to control for global chromatin accessibility^16,63^. H3K36me3 gene-body coverage was computed from annotated TSS to TTS, normalized by gene width and total fragment counts to account for sequencing depth and fragment-length distributions.

To compare with fragmentomic approaches, we used the publicly available implementation and recommended workflow from the EPIC-seq study (EPICSeqCode.v0.tar; https://epicseq.stanford.edu)^9^. Briefly, paired-end reads were trimmed using fastp v0.24.0^64^, aligned to hg19 using BWA-MEM v0.7.17 with extended insert-size modeling (-I 250,200,2000,10)^44^, and processed with samtools v1.15.1 to retain uniquely mapped fragments (MAPQ ≥ 1)^65^. Reads were sorted, deduplicated with Picard MarkDuplicates v2.6.0 (RRID:SCR_006525), and further filtered using samtools view -h -q 30 -F 3084, followed by restriction to canonical properly paired configurations (BAM flags 81, 83, 97, 99, 145, 147, 161, 163)^65^.

Fragmentomic features were computed using the publicly released EPIC-seq code from https://epicseq.stanford.edu^9^, including EPIC-seq-inferred expression, promoter fragmentation entropy (PFE), Gini index, orientation-aware fragmentation, Shannon index of fragment lengths, nucleosome depletion ratio (NDR), and GC content. One modification was applied to the PFE normalization step, informed by discussion with the original developers: the gamma-distribution shape parameter used in the pgamma (R stats v4.4.2) transformation was changed from 0.5 to 0.25 to better reflect entropy distributions in deeply sequenced cfDNA libraries and to avoid compression of values toward 1. All other components of the EPIC-seq pipeline were executed exactly as provided in the original code.

Because EPIC-seq uses Ensembl v75 TSS annotations, gene symbols were harmonized to GENCODE v27 using HGNChelper v0.8.15, yielding 17,348 shared genes for comparison^53^.

Predictive performance was assessed by computing gene-level Spearman correlations between each method’s inferred expression and matched tumor bulk RNA-seq (log₂(TPM+1)), using pairwise complete observations. Recovery of cancer-type-specific transcriptional programs was evaluated using the top 10 tumor-specific genes defined from matched bulk RNA-seq (see “Defining cancer- and normal-subtype-specific gene signatures”) and mean predicted expression of these gene sets was visualized as a 5×5 heatmap.

### 18. Evaluation of APEX and cfChIP-seq limit of SCLC detection through serial dilutions

#### 18.1. Serial dilution design and cfChIP-seq profiling

To evaluate whether APEX improves sensitivity for detecting tumor-derived chromatin signals beyond individual histone features, we performed a serial-dilution experiment using plasma from three patients with small-cell lung cancer (SCLC; tumor fractions 0.31, 0.41, and 0.55) and three non-cancer donors. Plasma cfDNA was extracted as described above under “Cell-free DNA isolation and low-pass whole genome sequencing (LP-WGS)”, and each SCLC plasma sample was diluted to match the cfDNA concentration of its paired non-cancer donor using dilution buffer (1% NP-40, 0.5% sodium deoxycholate, 0.1% SDS, protease inhibitor cocktail in PBS). To confirm dilution linearity, each sample was spiked with 192 pg of recombinant nucleosomes from the SNAP-ChIP® K-AcylStat™ Panel (EpiCypher #19-3100), including ∼8.3 pg of an H3K27ac nucleosome. Spike-in recovery was quantified following H3K27ac immunoprecipitation using qPCR on a LightCycler 480 with PowerUp™ SYBR™ Green Master Mix (Thermo Fisher Scientific, A25742; software version 1.5.1.62-SP3) and a universal forward primer (5′-CGTCGAACGCGCGATAT-3′) paired with an H3K27ac-specific reverse primer (5′-CGTATACGCGCGACA-3′).

For each SCLC–non-cancer pair, SCLC plasma was mixed with equimolar non-cancer plasma to generate a dilution series corresponding to the original tumor fraction multiplied by 1, 1⁄2, 1⁄4, 1⁄8, 1⁄16, 1⁄32, and 1⁄64 and a 0%-tumor (non-cancer-only) reference. Libraries were generated for all mixtures using the cfChIP-seq workflow described above under “Cell-free chromatin immunoprecipitation sequencing (cfChIP-seq)” and processed to obtain histone-mark coverage profiles and APEX-inferred gene-expression values. Although recombinant H3K27ac spike-in quantification demonstrated appropriate immunoprecipitation at all dilutions, endogenous H3K27ac yielded low total fragment numbers and enrichment scores and was therefore excluded from these analyses.

18.2. *Signature-level limit of detection analysis*

To estimate the analytical limit of detection, we quantified recovery of a SCLC-specific gene expression signature across the dilution series, using a previously published approach^14^. For each mixture, signal was computed using the top 100 genes from a predefined SCLC-specific geneset and normalized to a 500-gene WBC-specific background signature (*n* = 443 overlapping genes; see “Defining cancer- and normal-subtype-specific gene signatures”; **Supplementary Table 8**).

For each sample, APEX or histone coverage values were calculated for each SCLC gene and normalized for background WBC-derived signal by dividing by the mean signal across WBC-signature genes within the same sample. The resulting 100 normalized values per dilution condition were compared with the 0% non-cancer reference using a two-sided *t*-test, and the corresponding *t-*statistic was reported.

Three biological replicates were included per dilution, except for the 6.25% condition, which included two replicates. For each plasma metric, the relationship between dilution fraction (*x*-axis) and mean *t*-statistic (*y*-axis) was modeled using a logistic growth function implemented in GraphPad Prism (v10.4.1):

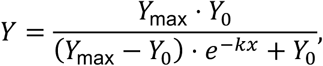

where *Y* is the predicted mean *t*-statistic at dilution fraction *x*, *Y*_max_ is the asymptotic maximum *t*-statistic, *Y*_0_ is the *t*-statistic at zero tumor fraction, and *k* is the growth rate constant. A *t-*statistic threshold of 2 (approximately corresponding to a two-sided *P* = 0.05) was selected as the detection cutoff.

#### 18.3. Single-gene-level limit of detection analysis

We also evaluated the analytical limit of single-gene detection across the top 100 genes from a predefined set of SCLC-associated genes, including *CHGA*, a lineage-specific marker for SCLC^26^, and *DLL3*, a clinically actionable target in SCLC^27^. For each dilution condition, gene-specific enrichment was calculated as signal relative to the non-cancer baseline using APEX-inferred expression, H3K4me3 promoter coverage, or H3K36me3 gene-body coverage.

To assess statistical significance, a null background distribution was generated by computing the same enrichment metric for all WBC-specific genes (*n* = 443). To avoid divisions by zero, a pseudocount of 1 was added to both the numerator and denominator. An empirical *P*-value were determined for each gene and dilution as the fraction of WBC-signature genes with enrichment equal to or exceeding that observed for the gene of interest. A gene was considered detected at a given dilution if enrichment relative to the WBC background exceed the WBC distribution and reached an empirical *P* < 0.05. For each gene, the limit of detection was defined as the lowest tumor fraction at which this criterion was met.

### 19. Benchmarking APEX–H3K4me3 against promoter coverage

To ask whether APEX applied to a single histone mark (APEX–H3K4me3) captures lineage-specific transcriptional information beyond raw H3K4me3 promoter coverage, we analyzed plasma cfChIP-seq data from a previously published study^16^. From this dataset, we selected 132 cfChIP-seq samples across colorectal cancer (*n* = 32), ER– breast cancer (*n* = 8), ER+ breast cancer (*n* = 6), hepatocellular carcinoma (*n* = 10), lung adenocarcinoma (*n* = 25), lung squamous cell carcinoma (*n* = 5), neuroendocrine prostate cancer (*n* = 9), and prostate adenocarcinoma (*n* = 37), applying the same strict criteria used in the original study, including H3K4me3 enrichment score >10, >2 million uniquely aligned fragments, and estimated tumor DNA fraction >0.03. We also removed patient-level duplicates and only included cancer subtypes with >3 biological replicates.

Cancer-subtype-specific marker genes were defined using bulk tumor RNA-seq profiles from the corresponding TCGA tumor types by computing a specificity score for each gene as described above (“Defining cancer- and normal-subtype-specific gene signatures”), retaining the top ten genes per cancer subtype. The union of these genes constituted the reference signature set. Plasma-derived APEX–H3K4me3 predicted expression values and raw H3K4me3 promoter coverage values (±3 kb around the TSS, normalized by DHS accessibility^16,63^) were subsetted to the intersecting marker set. For each cancer subtype, discriminatory performance was quantified by comparing the distribution of marker-gene signal in plasma samples from the cancer of interest versus all other cancers using a two-sided Wilcoxon rank-sum test, with –log₁₀(P) values. To visualize lineage-specific programs, we constructed marker-gene-by-cancer-subtype matrices in which each entry corresponds to the plasma signal of an individual signature gene (*n* = 10 per cancer subtype) evaluated across cancer subtypes. For visualization only, matrices were *z*-score normalized and values were smoothed using a centered rolling mean (window size = 3) along the cancer-subtype axis using the R zoo package (v1.8-15).

### 20. Classification of pancreatic basal and classical transcriptional states using APEX

To ask whether APEX-inferred gene expression more accurately discriminates basal versus classical transcriptional subtypes of pancreatic ductal adenocarcinoma (PDAC) than single-mark promoter or gene-body coverage, we applied PurIST, an established logistic-regression classifier trained on bulk tumor RNA-seq^28^. PurIST assigns PDAC samples to basal or classical subtypes using a predefined set of top-scoring gene pairs (TSPs) and is publicly available (https://github.com/naimurashid/PurIST; v02_12_2019^28^). We implemented the classifier using the published R code and released model object, applying the apply_classifier function with default settings to matched matrices of (i) bulk RNA-seq (reference), (ii) APEX-inferred gene expression, (iii) H3K4me3 promoter coverage, and (iv) H3K36me3 gene-body coverage.

Plasma cfChIP-seq profiles were obtained from an independent cohort of pancreatic cancer plasma samples from the PancSeq study (Semaan, Eid et al., manuscript in preparation, 2026; citation pending at time of submission) and approved for research under IRB-approved DFCI protocol 03-189. The cohort comprised samples collected at Dana-Farber Cancer Institute (DFCI; *n* = 26) and Gustave Roussy (*n* = 4). Only samples with matched H3K4me3 and H3K36me3 cfChIP-seq profiles and paired bulk tumor RNA-seq were included in this analysis (*n* = 8 basal; *n* = 22 classical). One sample (M2271) was also included in the MOSCATO–2 validation cohort and was analyzed consistently across datasets.

Subtype concordance between plasma-derived predictions and tumor-derived PurIST labels was assessed using two-sided Fisher’s exact tests. To quantify separation between predicted basal and classical states, PurIST-derived basal probabilities from each modality were compared using two-sided Wilcoxon rank-sum tests.

### 21. Metastatic bladder cancer plasma cohort and clinical annotation

#### 21.1. Patient cohort, sample collection, and clinical annotation

Patients with metastatic bladder cancer were identified from the Dana-Farber Cancer Institute (DFCI) Arthur and Linda Gelb Center for Translational Research biobank and associated clinical database under IRB-approved DFCI protocol 02-021. All samples and clinical data were de-identified prior to analysis.

Eligible patients had histologically confirmed urothelial carcinoma of the bladder with metastatic or unresectable disease and were treated with enfortumab vedotin (EV) monotherapy^66^. Plasma samples were required to be collected either within 90 days prior to initiation of EV, without interim cancer-directed therapy, or within 8 days following EV initiation (median [IQR], 0 [0–2] days); 5 out of 24 plasma samples were collected after treatment initiation (day 7 or 8). Only patients aged 18 years or older were included. Exclusion criteria included prior exposure to NECTIN4-targeting therapies and receipt of concurrent systemic anticancer therapies in combination with EV.

For each patient, a single aliquot of 810 μL biobanked plasma was obtained. Plasma samples had been processed and stored at −80 °C and had not undergone any prior freeze-thaw cycles before analysis.

Radiographic response assessments were performed by a trained clinician blinded to plasma-based analyses, in accordance with RECIST version 1.1 criteria. Best overall response was determined based on changes in the sum of target lesion measurements. Complete response (CR) was defined as disappearance of all target lesions, partial response (PR) as a ≥30% decrease in the sum of target lesion diameters, progressive disease (PD) as a ≥20% increase in the sum of target lesion diameters or appearance of a new target lesion, and stable disease (SD) as neither sufficient shrinkage to qualify for PR nor sufficient increase or new lesions to qualify for PD.

Clinical variables, including histologic diagnosis, treatment response, metastatic distribution (liver, brain, or bone), Karnofsky performance status (KPS), number of prior lines of therapy, and dose reductions prior to best response, were extracted from the electronic medical record (Epic Systems) by a trained clinician blinded to plasma-based analyses. Technical batch information for plasma processing was also recorded.

Plasma tumor fraction was inferred from cfDNA sequencing data using ichorCNA (v0.2.0)^21^. Copy-number profiles were generated genome-wide, and copy-number ratios (log₂ tumor/normal) were calculated for the *NECTIN4* locus using a 500 kb window spanning chr1:161–161.5 Mb (hg19). Default ichorCNA parameters were used, except for the use of 500 kb bins to enable resolution of focal copy-number alterations encompassing *NECTIN4*.

A de-identified summary of clinical and technical characteristics for the cohort (*n* = 24) is provided in **Supplementary Table 12**.

#### 21.2. Differential expression and survival analyses

Differential expression analysis of APEX-inferred gene expression profiles was performed using the limma R package (v3.62.2)^67^. Gene-level expression estimates were modeled using linear models with group-specific coefficients, enabling direct estimation of expression differences between predefined comparison groups (for example, responders versus non-responders to enfortumab vedotin; **Supplementary Table 13**). Empirical Bayes moderation was applied to stabilize variance estimates, and log₂ fold changes and associated *P*-values were computed for all genes.

Survival analyses were performed using the survival R package (v3.8.3)^68^. For univariate analyses, patients were stratified into high and low groups based on median dichotomization of the feature of interest. Progression-free survival and overall survival were evaluated using Kaplan-Meier estimates and compared between groups using two-sided log-rank tests. Hazard ratios (HRs) and 95% confidence intervals (CIs) were estimated using Cox proportional hazards models.

Multivariable Cox proportional hazards models were used to assess whether plasma-inferred molecular features were independently associated with survival outcomes after adjustment for relevant clinical covariates. Candidate covariates were first screened in univariate Cox models, and variables with the strongest univariate associations were prioritized to limit model complexity. Based on this screening, APEX-inferred *NECTIN4* expression, presence of visceral metastases (liver and/or brain), and plasma tumor fraction (estimated using ichorCNA v0.2.0^21^) were included in the final multivariable models. Statistical significance was assessed using Wald tests, and forest plots were generated using the forestmodel R package (v0.6.2). All statistical tests were two-sided.

### 22. Immunohistochemistry and pathologic scoring

Tumor samples selected for DLL3 and TROP2 immunohistochemistry (IHC) were chosen based on plasma-inferred expression levels derived from APEX, with cases dichotomized into high and low groups using a median cutoff and representative samples taken from the highest and lowest inferred expression values. Sample selection was otherwise restricted only by the availability of matched tumor tissue and the suitability of formalin-fixed paraffin-embedded (FFPE) material for immunohistochemical staining. Pathologic evaluation was performed blinded to plasma-derived expression estimates.

IHC was performed on 3-µm–thick deparaffinized FFPE tissue sections using an automated staining platform (Ventana Benchmark; Ventana Medical Systems, Tucson, AZ, USA) at the PETRA platform, Gustave Roussy. Serial sections were stained with antibodies against DLL3 (clone SP347, prediluted; Roche, France; 08416931001) and TROP2 (clone EPR20043, 1:500; Abcam, USA; ab214488). Stained slides were digitized at ×20 magnification using a Slideview VS200 scanner (Olympus, Tokyo, Japan), ensuring high spatial resolution for evaluation.

DLL3- and TROP2-stained slides were evaluated by a pathologist using standard light microscopy. Cases with major tissue artifacts in the tumor area that precluded reliable evaluation were excluded from analysis. Scoring was performed at ×20 magnification in malignant cells using a semi-quantitative approach incorporating both the percentage of positive tumor cells (0–100%) and staining intensity (0, none; 1+, weak; 2+, moderate; 3+, strong). An H-score was calculated as

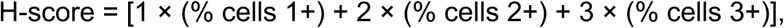

yielding a score range from 0 to 300. Any immunoreactivity below 1+ intensity was considered background or non-specific (score 0). DLL3 and TROP2 expression was considered positive when ≥1% of tumor cells demonstrated cytoplasmic and/or membranous staining at any intensity.

### 23. APEX-based nomination of expression-based cancer targets

Expression-based therapeutic targets were curated from the U.S. Food and Drug Administration (FDA) website and from a published compendium of cancer cell-surface protein targets^34^. FDA-approved targets were restricted to indications in solid tumors to minimize prioritization of hematopoietic markers arising from background blood signal. *CLDN18* was excluded because the approved antibody zolbetuximab specifically targets the *CLDN18.2* isoform, which cannot be resolved by our existing approach^69^.

For the expanded analysis, the FDA-approved target list was combined with candidate cancer therapeutic targets from “*Supplementary Table 9: List of the cancer-specific GESPs (caGESPs)*”, excluding targets specific to acute myeloid leukemia^34^. Genes overlapping with hematopoietic genes, which were defined as those with higher mean expression in blood-and immune- dominant tissues in GTEx (e.g. whole blood, EBV-transformed lymphocytes, and spleen) than in other samples, were removed, leaving 291 non-hematopoietic cancer-specific cell surface targets (**Supplementary Table 8**).

The pan-solid cancer cohort comprised all plasma samples included in this study from the MOSCATO–1 and MOSCATO–2 clinical trial cohorts, the pancreatic cancer cohort, the metastatic bladder cancer cohort, and the previously published cfChIP-seq cohort^16^. Only one sample per patient was included, selected at random among replicates, with no minimum tumor fraction threshold applied. Samples from the Baca *et al., Nat. Med.*, 2023 cohort^16^ were included if they met a permissive quality criterion of enrichment score >7.

Cancer types were summarized into unified categories as follows: Breast (breast cancer); Colorectal (colorectal cancer); Upper gastrointestinal (esophageal adenocarcinoma, esophageal squamous cell carcinoma, small intestine adenocarcinoma, and gastric cancer); Pancreatobiliary (pancreatic cancer, cholangiocarcinoma); Gynecologic (endometrial and ovarian cancer); Liver (hepatocellular carcinoma); NSCLC (lung adenocarcinoma, lung squamous cell carcinoma); SCLC / Neuroendocrine (small-cell lung cancer, neuroendocrine lung tumors, small-cell bladder cancer, Merkel cell carcinoma); Prostate / NEPC (castration-sensitive, castration-resistant, and neuroendocrine prostate cancer); Urothelial (urothelial carcinoma); Central nervous system (glioma); and Other (vesicular adenocarcinoma, melanoma, renal cell carcinoma, mesothelioma, myoepithelial carcinoma, medullary thyroid cancer, head and neck cancer, otherwise unspecified).

APEX inference was performed using H3K4me3 alone when H3K36me3 was unavailable, and using both marks when available, with default model parameters. In total, 417 plasma samples yielded APEX-inferred gene expression values for this analysis. For each target gene, high expression was defined as inferred expression greater than or equal to the 90th percentile of that gene’s inferred expression across the cohort. Thresholding was applied independently for each gene.

### 24. Statistical analyses and reproducibility

No statistical methods were used to predetermine sample size. Cohort sizes were dictated by the availability of matched plasma and tumor material. Measurements were obtained from distinct biological samples (one patient per sample) unless otherwise indicated. For clinical cohorts, investigators performing plasma-based analyses were blinded to matched tumor reference data and clinical outcomes at the time of analysis. All statistical tests were two-sided unless otherwise noted, and a *P*-value < 0.05 was considered statistically significant. Associations between plasma-derived features and tumor RNA-seq were assessed using Spearman correlation, group comparisons using Wilcoxon rank-sum, *t*-test, or Fisher’s exact tests, differential expression using limma with empirical Bayes moderation, and survival outcomes using Kaplan-Meier analysis and Cox proportional hazards models. Machine-learning models were evaluated using leave-one-out cross-validation and independent validation cohorts, with performance assessed by Spearman correlation and root mean squared error. Analyses were performed in R (v4.4.2) using publicly available software with versions specified in the **Methods**. For routine plotting and data manipulation, we used the R packages ggplot2 (v4.0.1), data.table (v1.17.8), and dplyr (v1.1.4).

### 25. Data availability

Raw human sequencing data generated and analyzed in this study are being deposited in dbGaP under controlled access to protect participant privacy (accession pending). These datasets include plasma cfChIP-seq profiling of histone modifications (H3K4me3, H3K36me3, and H3K27ac), cfMeDIP-seq, low-pass and deep whole-genome sequencing, matched tumor RNA-seq where applicable, and associated clinical metadata, as described in the Methods. Data are being deposited from the MOSCATO training cohort (*n* = 15 patients), validation cohorts (*n* = 34 patients), an external cohort (*n* = 2 patients), limit-of-detection dilution series experiments (*n* = 6 patients), the DFCI metastatic bladder cancer cohort (*n* = 24 patients), and the DFCI/Gustave Roussy metastatic pancreatic cancer cohort (*n* = 30 patients). Plasma H3K4me3 cfChIP-seq data with matched phenotype annotations from a previously published cohort were also used, with data availability as described in that study^16^.

Processed data from these cohorts, including bigWig coverage tracks and BED-like fragment files, have been deposited in Gene Expression Omnibus (GEO; accession GSE318228). APEX-inferred gene expression profiles for all cohorts are provided as **Supplementary Data**. In addition, raw and processed data from gene-body cfChIP-seq benchmarking analyses using mixed samples derived from prostate cancer cell lines and non-cancer donors (*n* = 84 epigenomic profiles), as well as LNCaP chromatin fragmentation comparisons between MNase- and Covaris-based protocols (*n* = 10 biological replicates per condition), have also been deposited in GEO (accession GSE318228).

Raw bulk tumor RNA-seq data for patients enrolled in the MOSCATO–1 and MOSCATO–2 trials are available through the European Genome–Phenome Archive (EGA; accession EGAC00001002937). Raw and processed bulk tumor RNA-seq data for the metastatic pancreatic cancer cohort were obtained from a companion study, as described in the **Methods**, and are available through their corresponding repository (Semaan, Eid et al., manuscript in preparation, 2026; citation and accession pending at time of submission). Processed bulk RNA-seq expression matrices used in this study are provided as **Supplementary Data**.

Publicly available datasets used for model training, benchmarking, or comparative analyses include bulk RNA-seq expression and matched clinical data from TCGA and GTEx accessed via the UCSC Xena Browser (https://xenabrowser.net/datapages/?cohort=TCGA%20TARGET%20GTEx; accessed June 7, 2025) and the TCGABiolinks R package (v2.36.0)^50^; epigenomic and transcriptomic reference datasets from the Roadmap Epigenomics Project (https://egg2.wustl.edu/roadmap/web_portal/processed_data.html); small-cell lung cancer RNA-seq data from George et al. (Nature, 2015; PMID: 26168399; Supplementary Table 10)^51^; metastatic prostate cancer RNA-seq datasets from the Prostate Cancer Atlas, including the “WCDT-MCRPC” and “T.C.B. 2015” studies (https://www.prostatecanceratlas.org/api/files/public/gc_txi.abundance.fst; dbGaP accession phs000909.v1.p1); and control LNCaP RNA-seq data from GEO (GSE128749). A curated list of candidate cancer cell-surface targets was obtained from Hu et al. (Nature Cancer, 2021; PMID: 35121907; Supplementary Table 9)^34^.

Public genomic and epigenomic annotations were obtained from ENCODE genomic exclusion regions (ENCFF001TDO; https://www.encodeproject.org/files/ENCFF001TDO/@@download/ENCFF001TDO.bed.gz), the Boyle Lab hg19 blacklist (https://github.com/Boyle-Lab/Blacklist/blob/master/lists/hg19-blacklist.v2.bed.gz), reference genome coordinates from the BSgenome.Hsapiens.UCSC.hg19 R package (v1.4.3), protein-coding gene annotations from GENCODE (GRCh37, release 27), enhancer annotations from GeneHancer (v5.22, GRCh37)^22^, DNase I hypersensitive site annotations (https://zenodo.org/record/3838751/files/DHS_Index_and_Vocabulary_hg19_WM20190703. txt.gz), genome-wide ChromHMM 18-state annotations across 833 biosamples from the Epimap project (hg19; https://personal.broadinstitute.org/cboix/epimap/ChromHMM/observed_aux_18_hg19/CALL S/), and public ChIP-seq and paired RNA-seq datasets from the Roadmap Epigenomics Project (https://egg2.wustl.edu/roadmap/web_portal/processed_data.html).

### 26. Code availability

All source code and trained models required to reproduce the analyses in this study will be made publicly available upon publication. Preprocessing of cfChIP-seq data was performed using the Nextflow-based SNAPIE pipeline (v1.5.1), available at https://github.com/prc992/SNAPIE^43^. EPIC-seq data were processed using the EPIC-seq framework (https://epic.stanford.edu)^9^. APEX (v4.0.0) was developed in this study and is distributed as a fully documented R package providing functions for cfChIP-seq quality assessment, extraction of epigenomic and fragmentomic feature, inference of genome-wide gene expression using pretrained models, differential expression and gene set analyses, and data visualization, including interactive plots. The software is packaged as a fully reproducible Docker container that encapsulates all dependencies and workflows, and includes comprehensive documentation, vignettes, example inputs, and a Snakemake^70^ workflow for scalable execution on HPC systems. During peer review, access to the APEX source code, trained models, and Docker container is provided to reviewers via a tarball of the corresponding GitHub repository. Upon publication, the source code, container, trained model files, and code to generate figure and analyses in the manuscript will be made publicly accessible for non-profit academic use via GitHub.

## Supplementary Note

### Accounting for variable promoter usage across tumor types

Promoter activity is highly tissue- and lineage-specific, and many genes use alternative promoters depending on cellular context^25^. In our cohort, 29% of genes exhibited H3K4me3 signal peaks at locations distinct from the GENCODE v27-annotated TSS, and 64% of these varied between the 15 evaluated samples (**Supplementary Table 5**). To address this heterogeneity in promoter usage, we developed a method to dynamically reassign each gene’s TSS per sample based on the position of maximal H3K4me3 signal among annotated TSSs (**Methods**, “Development of APEX: Selection of alternate promoter”). Genes for which TSS reassignment left a residual gene body shorter than 500 bp were excluded from downstream analyses, as the resulting bins were too small for robust feature estimation. Overall, this adjustment significantly improved expression correlations for narrow histone marks such as H3K4me3 and H3K27ac, particularly for context-dependent genes like *ESR1*, but had less impact on broad marks such as H3K36me3 (**Extended Data Fig. 5**).

### Selection of APEX model

To identify a parsimonious APEX model that preserved maximal predictive performance while minimizing redundant information and computational cost, we systematically evaluated the contribution of individual genomic regions, histone marks, and feature groups using the training cohort (*n* = 15 plasma–tumor pairs). First, we assessed the contribution of each histone modification by excluding or isolating H3K4me3, H3K27ac, and H3K36me3 features (**Extended Data Fig. 6a**). Omitting H3K27ac had no measurable effect on error, and models using H3K27ac alone performed worst, indicating limited independent value of this mark in the cfChIP-seq setting. In contrast, both H3K4me3 and H3K36me3 individually reduced performance when excluded, and neither mark alone achieved maximal predictive accuracy, demonstrating complementary and non-redundant information content.

Next, we quantified the effect of excluding each genomic region (e.g. promoters, gene bodies, intergenic enhancers, and intragenic enhancers) on model error using leave-one-region-out cross-validation. Removing intergenic or intragenic enhancers did not significantly alter performance (**Extended Data Fig. 6b**), whereas promoter and gene-body regions were consistently required for optimal accuracy.

We further tested whether adding 5-methylcytosine-based fragment enrichment (cfMeDIP-seq) improved performance relative to cfChIP-based models alone. Across an independent cohort (*n* = 18), inclusion of methylation features did not significantly reduce RMSE (**Extended Data Fig. 6c**), suggesting minimal added benefit beyond chromatin-encoded signals in this context.

To evaluate redundancy across broader feature classes, we performed leave-one-feature-group-out and leave-all-but-one-feature-group analyses across fragment-based and chromatin-based features (4-mer end-motifs, GC-weighted coverage, fragment length metrics, and histone mark coverage). No single group accounted for the full model performance and removing any one class did not significantly degrade accuracy (**Extended Data Fig. 6d**), indicating that information is distributed across features and largely orthogonal. Based on this, we retained both chromatin-derived and fragmentomic features, reasoning that orthogonal signals are advantageous when working with sparse cfDNA data.

From these analyses, we defined a refined APEX model that included only the two essential histone marks (H3K4me3 and H3K36me3) and only the promoter and gene-body regions, reducing the feature set from 3,573 features in the full model to 1,532 features (**Fig. 6e**). APEX variants using a single histone mark (766 features) were also evaluated for parsimony and operational simplicity.

Finally, we benchmarked runtime performance across all models using the training cohort. All tests were performed on an Apple M3 Max MacBook Pro (16-core CPU, 64 GB RAM). With optimized XGBoost parameters (maximum depth = 7, minimum loss reduction = 1.0, feature fraction = 0.7, learning rate = 0.10, sample fraction = 1.0), the full model required a median of 10.5 minutes per sample, whereas the refined model required 6.5 minutes per sample without any loss of accuracy (**Extended Data Fig. 6g**). Single-mark APEX variants using only H3K4me3 or only H3K36me3 further reduced runtime (median 2.5 and 4.2 minutes per sample, respectively) but at the cost of reduced predictive accuracy (**Extended Data Fig. 6f,g**).

Based on the combined criteria of predictive performance, non-redundancy, feature complementarity, and runtime efficiency, we selected the refined APEX model, which incorporates H3K4me3 and H3K36me3 promoter and gene-body features, as the preferred model for downstream analysis (**Extended Data Fig. 6h**).

